# *Anopheles* salivary apyrase regulates blood meal hemostasis and drives malaria parasite transmission

**DOI:** 10.1101/2023.05.22.541827

**Authors:** Zarna Rajeshkumar Pala, Thiago Luiz Alves e Silva, Mahnaz Minai, Benjamin Crews, Eduardo Patino-Martinez, Carmelo Carmona-Rivera, Paola Carolina Valenzuela-Leon, Ines Martin-Martin, Yevel Flores-Garcia, Raul E. Cachau, Naman Srivastava, Ian N. Moore, Derron A. Alves, Mariana J Kaplan, Elizabeth Fischer, Eric Calvo, Joel Vega-Rodriguez

## Abstract

Mosquito salivary proteins play a crucial role in regulating hemostatic responses at the bite site during blood feeding. In this study, we investigate the function of *Anopheles gambiae* salivary apyrase (AgApyrase) in *Plasmodium* transmission. Our results demonstrate that salivary apyrase interacts with and activates tissue plasminogen activator, facilitating the conversion of plasminogen to plasmin, a human protein previously shown to be required for *Plasmodium* transmission. Microscopy imaging shows that mosquitoes ingest a substantial amount of apyrase during blood feeding which reduces coagulation in the blood meal by enhancing fibrin degradation and inhibiting platelet aggregation. Supplementation of *Plasmodium* infected blood with apyrase significantly enhanced *Plasmodium* infection in the mosquito midgut. In contrast, AgApyrase immunization inhibited *Plasmodium* mosquito infection and sporozoite transmission. This study highlights a pivotal role for mosquito salivary apyrase for regulation of hemostasis in the mosquito blood meal and for *Plasmodium* transmission to mosquitoes and to the mammal host, underscoring the potential for new strategies to prevent malaria transmission.

## Introduction

Malaria, a debilitating and sometimes fatal infectious disease, continues to pose a major public health challenge, with approximately 247 million cases and 619,000 deaths globally in 2021 [1]. The transmission of the causative agent, *Plasmodium* parasite, occurs via the bite of an infected *Anopheles* mosquito. The parasite undergoes sexual reproduction and differentiates into male and female gametes inside the mosquito midgut after ingesting a blood meal from an infected human, producing motile ookinetes that invade the midgut epithelium to form oocysts that produce the infectious sporozoites [2]. The development of the parasite in the mosquito midgut and transmission of sporozoites to humans represent critical bottlenecks in the life cycle, making them attractive targets for the development of new malaria interventions. Recent studies have shown that *Plasmodium* co-opts host proteins, such as the serine protease plasmin, to facilitate infection in both the mosquito vector and the human host [3–5].

Fibrinolysis, a crucial process for maintaining homeostasis in mammals, involves the degradation of fibrin clots by plasmin. This process is initiated by the activation of plasminogen into plasmin by plasminogen activators, such as tissue-type (tPA) and urokinase-type (uPA) plasminogen activators. In a positive feedback loop, the basal activity of tPA and uPA activates plasminogen into plasmin, and plasmin further activates tPA and uPA thereby, enhancing the overall plasminogen activation [6]. We recently demonstrated that *Plasmodium* gametes hijack tPA to activate plasminogen on their surface, allowing it to degrade fibrin in the blood bolus and facilitate gamete migration [3]. Moreover, we showed that sporozoites also hijack plasmin for degradation of ECM proteins, which facilitates its motility in the skin and liver [3].

Mosquito saliva contains numerous proteins with anti-hemostatic and immunomodulatory functions that not only play a role in facilitating blood feeding, but also impact pathogen transmission [7–11]. While the role of mosquito saliva and some salivary proteins in *Plasmodium* sporozoite transmission has been studied, our understanding of their impact on *Plasmodium* transmission from humans to mosquitoes, and from mosquitoes to humans, remains limited. Recent studies showed that salivary saglin and SGS1 ingested during blood feeding impacts *Plasmodium* midgut infection however, the mechanism of action is still unknown [12, 13]. In this study, we provide the mechanism by which mosquito salivary apyrase, ingested during blood feeding, enhances fibrinolysis and inhibits platelet activation in the mosquito blood bolus, which facilitates *Plasmodium* infection in the mosquito vector and transmission to the mammalian host.

## Results and Discussion

To evaluate the impact of mosquito salivary proteins on the fibrinolytic system, we investigated the ability of *Anopheles gambiae* salivary gland protein extracts and saliva to activate single-chain tissue plasminogen activator (sc-tPA). Using a tPA fluorogenic substrate (D-Val-Leu-Lys-7-amido-4-methylcoumarin), we observed increased tPA activity in response to both salivary gland extracts (Fig. 1A) or saliva (Fig. S1A). Activation of tPA was lost upon heating the salivary glands to 65 or 100 °C (Fig. 1A), suggesting a salivary protein as the tPA activator. To identify the putative salivary tPA activator, we fractionated salivary proteins by size-exclusion chromatography and identified fraction Z8 as the strongest tPA activator, whereas the adjacent fractions Z7 and Z9 activated tPA at lower levels (Fig. 1B). Mass spectrometry analysis of these fractions identified a total of 152 unique proteins (Table S1), which were shortlisted to eight potential tPA activators (Table S2 and Fig. S2A) based on the presence of secretion signal, presence in adjacent fractions, and absence from fractions A5 and B3 which did not activate tPA (Table S1). The eight candidates were expressed in HEK293 cells and purified (Fig. S2B), and only AGAP011026-PA, annotated as a 5’ nucleotidase ecto (Ag5’NTE) in VectorBase, activated tPA in the fluorogenic assay at levels comparable to saliva (Fig. 1C and Fig. S2C), identifying salivary *An. gambiae* 5’ NTE as a tPA activator. In mammals, activation of single-chain tPA (sc-tPA) takes place by proteolytic cleavage by plasmin generating two-chain tPA (tc-tPA), or by allosteric interaction with other proteins like fibrin [14, 15]. ELISA overlay assays showed that recombinant Ag5’NTE (rAg5’NTE) directly interacted with tPA (Fig. 1D) but did not enhance cleavage of sc-tPA into two-chain tPA (tc-tPA) (Fig. S1B), suggesting an allosteric activation mechanism. An unrelated protein amongst the eight candidate proteins, (AGAP007393) did not interact with tPA (Fig. 1D). Next, using a colorimetric assay, we tested whether the rAg5’NTE-activated tPA in turn activated plasminogen to plasmin [3]. rAg5’NTE-activated tPA resulted in the highest activation of plasminogen to plasmin when compared to the positive controls of plasmin, or plasminogen activation by sc-tPA or two-chain tPA (tc-tPA) (Fig. 1E). These results demonstrate the important role of Ag5’NTE in the activation of fibrinolytic system. Based on our previous report showing that the fibrinolytic system enhances parasite infection in the mosquito and transmission to the mammalian host [3], we hypothesized that Ag5’NTE may facilitate *Plasmodium* transmission by activating fibrinolysis.

**Figure 1:**
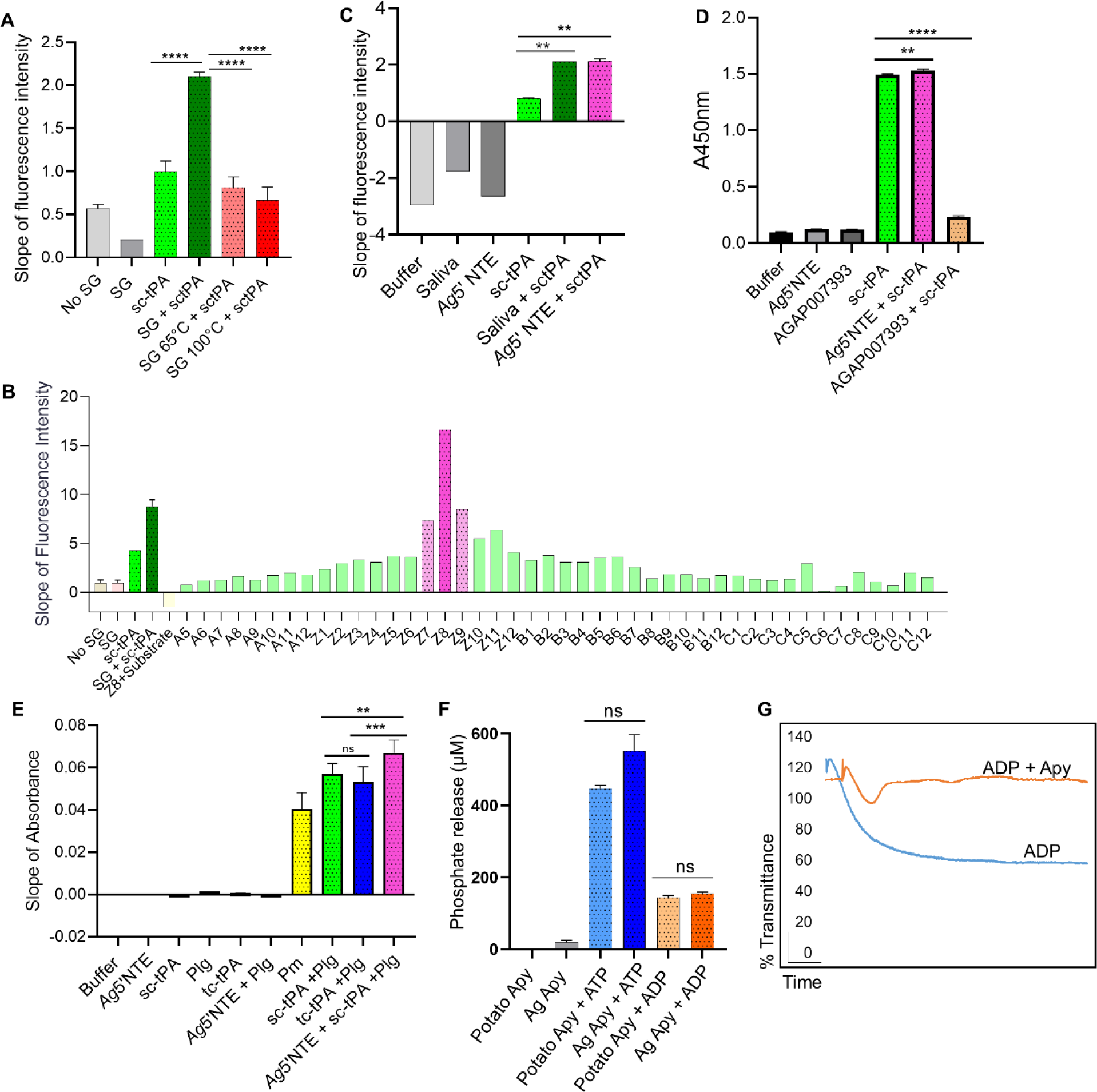
*An. gambiae* salivary apyrase activates tPA. (**A**) Fluorogenic assay for single-chain tPA (sc-tPA) activation reveal that mosquito salivary gland extracts (SG) activate sc-tPA. Heating the extract at 65°C or 100°C prevents tPA activation. ****P <0.0001. (**B**) Fractions from *An. gambiae* salivary gland extracts obtained by size-exclusion chromatography were tested for tPA activation. Fraction Z8 showed higher sc-tPA activation compared to sc-tPA or SG alone. (**C**) *An. gambiae* salivary 5’ nucleotidase ecto (Ag5’NTE) identified as the saliva tPA activator. Ag5’NTE activates tPA at similar levels as *An. gambiae* saliva. **P <0.01. (**D**) Interaction of tPA with Ag5’NTE using ELISA overlays. rAg5’NTE and an unrelated protein (AGAP007393) were coated on the plate. Wells were incubated with or without sc-tPA. Anti-tPA antibodies were used to detect sc-tPA. Ag5’NTE interact with sc-tPA, whereas AGAP007393 does not interact (mustard yellow). A well was coated with sc-tPA as positive control. **P <0.01; ****P <0.0001. (**E**) Plasminogen activation in the presence of rAg5’ NTE and tPA measured by a colorimetric assay. Ag5’NTE activated tPA in-turn activates plasminogen to plasmin (dark pink) at higher levels than plasmin, sc-TPA or tc-tPA alone. **P <0.01; ****P <0.001; ns: not significant. (**F**) Ag5’NTE (AGAP011026) is an apyrase. Inorganic phosphate released was measured using Fiske Subbarow reagent. Ag5’NTE (AGAP011026), now referred as AgApyrase, released inorganic phosphate from ATP and ADP when compared to the positive control potato apyrase. ns: not significant. (**G**) rAgApyrase inhibits ADP-mediated platelet aggregation. Platelet-rich plasma was incubated with the platelet aggregation agonist ADP and platelet aggregation was measured by light transmittance over 6 min. rAgApyrase inhibited the platelet aggregation induced by ADP (orange) in contrast to the control buffer with ADP (blue) which aggregated all the available platelets. Data from three (A, D, E, F-ATP), two (C, F-ADP) or one (B) independent experiments. Groups were compared with an ordinary one-way ANOVA followed by Sidak’s multiple comparison test for pairwise comparison (A, C, D) or paired two-tailed t-test (E, F).

The 5’ nucleotidase ecto is an extracellular protein, normally attached to the membrane by a glycosyl phosphoinositol (GPI) anchor or secreted, that hydrolyzes AMP to adenosine and phosphate [16]. However, AGAP011026-PA sequence does not have the predictive site for a GPI anchor indicating it is secreted [17, 18]. *Anopheles* mosquitoes express two 5’ nucleosidases (AGAP011971 and AGAP011026) in the salivary glands and based on their expression pattern, AGAP011971 was classified as a real apyrase and AGAP011026 as a 5’ nucleosidase ecto however, no biochemical assays has been perform to validate their activity [19]. Using a biochemical colorimetric assay to measure phosphate release as an indicator of AMP hydrolysis [20], we found that rAg5’NTE did not show any activity, whereas a strong activity was observed in the positive control with human 5’ NTE (CD73) (Fig. S3A). 5’ nucleotidase ecto is a member of the 5’-nucleotidase family that also include apyrases, which hydrolyzes ATP and ADP to AMP in the presence of a bivalent metal ion [21, 22]. Upon incubation of rAg5’NTE with either ATP or ADP in the presence of calcium, we observed significant release of inorganic phosphate at levels similar to the positive control potato apyrase (Fig. 1F and Fig. S3B). Apyrases belong to three families: the 5’ nucleotidases family which includes apyrases from *Aedes aegypti* [23], *An. gambiae* [24] and *Triatoma infestans* [25]; and a novel apyrase family which includes apyrases from *Cimex lectularius* [26], *Phlebotomus papatasi* [27], *Lutziomyia longipalpus* [28], *Drosophila melanogaster* [28], and humans [29], and the CD39 family, which includes apyrases of humans [30], and possibly, fleas [31]. The 5’ nucleotidase type functions in the presence of Ca^2+^ or Mg^2+^, and the *Cimex* type works only with Ca^2+^ [32]. Strikingly, the rAg5’NTE hydrolyzed ADP like a *Cimex*-type apyrase in the presence of Ca^2+^ only (Fig. S3C) despite being phylogenetically distant (Fig S4A). The amino acid residues binding to calcium were well conserved amongst the members belonging to both 5’ nucleotidase type and *Cimex* type apyrases (Fig. S4B). Salivary apyrases from hematophagous insects inhibit ADP-mediated platelet activation and aggregation which could have significant implications in human hemostatic responses during insect feeding [33]. We observed that rAg5’NTE (AGAP011026-PA) strongly inhibits ADP-mediated platelet aggregation *in vitro* (Fig. 1G). These results confirm that AGAP011026-PA (Ag5’NTE) is the *An. gambiae* salivary apyrase, henceforth referred to as AgApyrase (AgApy). In silico structural analysis of the interaction between AgApyrase and human tPA produced a complex model with a buried surface area 1467 A^2^, where the interaction between the two molecules is mediated by one of the tPA protruding loops in the tPA serine protease domain (Fig. S5A). The AgApyrase residues mediating the interaction are mostly located in the nucleotide binding site and the nucleosidase catalytic site. It is worth noticing that the residues involved in the Apyrase-tPA contact are highly conserved across the selected set (Fig. S5B). Further biochemical and structural analysis needs to be performed to validate the predicted interaction.

Our study builds on our previous finding that fibrin polymerization in the mosquito midgut increases the viscosity of the blood meal, hampering parasite motility and infectivity [3]. Since we showed that salivary AgApyrase activates the fibrinolytic system (Fig. 1E), we postulate that ingestion of AgApyrase enhances fibrin degradation, thereby boosting parasite infection in the mosquito midgut. To confirm the presence of salivary apyrase in the mosquito blood bolus, we performed immunohistochemistry with anti-apyrase antibodies on blood (human)-fed mosquito midguts dissected 30 min post feeding and unfed whole mosquitoes. We observed that *An. gambiae* mosquitoes ingest a significant amount of apyrase during feeding (Fig. 2A and Fig. S6A and S6B), which is supported by other studies showing the ingestion of saliva during blood feeding [4, 12, 34]. The intensity of apyrase staining in the blood bolus shows that the mosquito ingests a substantial amount of saliva while feeding on blood, which has important implications for its effect in modulating hemostatic and inflammatory responses in the mosquito blood bolus. Apyrase could be detected in the lumen of unfed mosquitoes (Fig. S6C), presumably from saliva ingested without feeding on blood since this apyrase is not expressed in mosquito midguts according to published transcriptomes included in VectorBase [35–37]. Western blot analysis showed the specificity of the anti-AgApyrase antibodies which only reacted with the homogenate of one blood fed mosquito midgut and not with human plasma or serum (Fig. S6D). These results confirm that, when taking a blood meal, a large amount of salivary AgApyrase is ingested by the mosquito.

**Figure 2:**
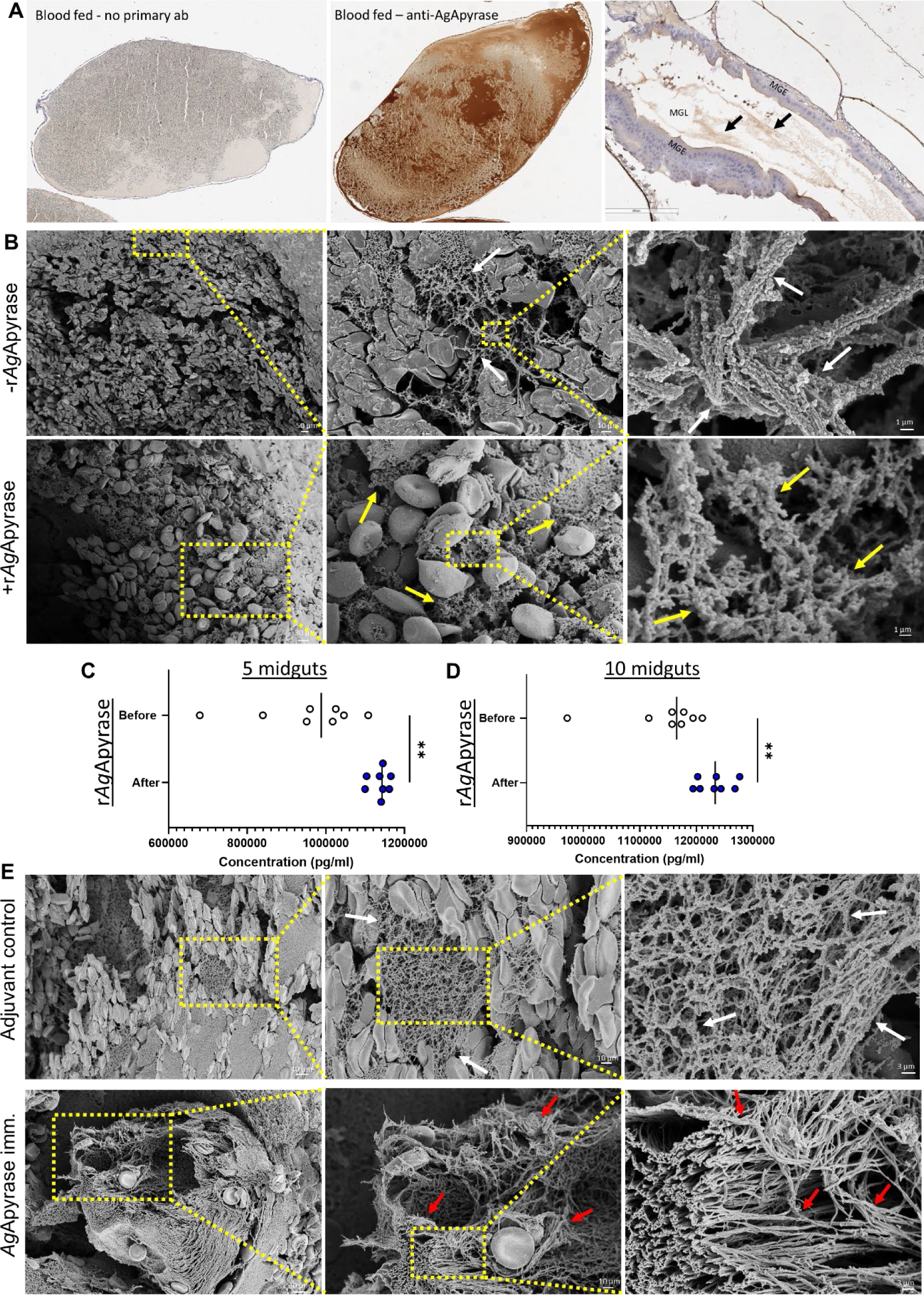
Salivary AgApyrase is ingested during blood feeding and enhances fibrinolysis in the blood bolus. (**A**) Immunohistochemistry performed on blood fed mosquito midgut shows ingestion of salivary AgApyrase in the mosquito midgut (dark brown patches in the middle panel) by staining with anti-apyrase antibodies. Blood fed mosquito midgut with no primary antibody did not show any signal (absence of brown patches in the first panel). Unfed mosquito midgut stained with anti-apyrase antibodies showed faint brown patches (black arrows in the third panel). MGE: midgut epithelium, MGL: midgut lumen. (**B**) Scanning electron microscopy (SEM) of blood boluses before and after supplementation with rAgApyrase. *An. gambiae* female mosquitoes were fed on mice before or after intravenous injection of rAgApyrase. Midguts were dissected at 30 min (or at 4 h shown in Fig. S8) post feeding. Note the well organize fibers formed in the blood bolus before the supplementation with rAgApyrase (white arrows) as compared to the thinner and sponge-like structure formed after rAgApyrase supplementation (yellow arrows). (**C, D**) Effect of rAgApyrase on D-dimer formation in blood boluses. Mosquitoes were fed on mice before or after intravenous injection of rAgApyrase, and midguts were dissected 30 min post feeding to measure D-dimer formation. Assays were done with pools of 5 (C) or 10 (D) midguts. Supplementation of the blood meal with rAgApyrase increases D-dimer formation. Data pooled from two independent experiments. Unpaired t-test, **P <0.002. (**E**) SEM of blood boluses from mosquitoes fed on rAgApyrase immunized mice. Control mosquitoes were fed on mice treated with adjuvant. Midguts were dissected 30 min post feeding. Note the increase in fiber organization and thickness (red arrows) observed in boluses from mosquitoes that fed on rAgApyrase immunized mice as compared to adjuvant treated mice.

Next, we tested whether AgApyrase enhanced fibrinolysis in the midgut. Using scanning electron microscopy (SEM), we compared the formation of fibrin networks in the midgut blood bolus of mosquitoes that fed on mice before and after intravenous injection of rAgApyrase. Dissection at 30 minutes or 4 hours post-feeding showed that midguts from control mosquitoes (before apyrase injection) exhibited a thick and organized fibrin polymer network, whereas those fed on rAgApyrase injected mice displayed a looser sponge-like mesh (Fig. 2B and Figs. S7 and S8). Immune transmission electron microscopy of mosquito blood boluses confirmed the presence of fibrinogen in the space between red blood cells (Fig. S9), supporting that the observed fibers in transmission electron microscopy are fibrin. To determine whether we observe an increase in fibrinolysis induced by apyrase ingestion, we measured the concentration of D-dimers, degradation products of cross-linked fibrin, formed in the midgut blood bolus of mosquitoes fed on mice before and after rAgApyrase injection. Results from a competitive ELISA assay showed a significant increase in D-dimers in mosquitos fed on rAgApyrase supplemented mice, showing that apyrase ingestion enhanced fibrinolysis in the blood bolus (Fig. 2C and 2D). Further, SEM of blood boluses from mosquitoes fed on rAgApyrase immunized mice also indicated a more organized fibrin polymerization when compared to mosquitoes fed on the adjuvant control group (Fig. 2E and Fig S10). In conclusion, our results demonstrate that salivary apyrase ingested by *An. gambiae* mosquitoes during blood feeding increases fibrinolysis in the midgut, leading to the formation of a looser fibrin polymer network in the blood bolus.

Platelets and fibrin play crucial roles in coagulation. Platelets initiate the formation of a primary platelet plug at injury sites, whereas fibrin stabilizes it [38]. The release of ADP from activated platelets during coagulation further amplifies platelet aggregation [39]. Some hematophagous arthropods such as mosquitoes can inhibit ADP-mediated platelet aggregation through their salivary apyrase [40]. Our study demonstrated that recombinant apyrase from *An. gambiae* mosquitoes (rAgApyrase) inhibits ADP-mediated platelet aggregation *in vitro* (Figure 1G). To investigate the effect of salivary apyrase on platelet activation and aggregation in the mosquito midgut, we performed immunohistochemistry on midgut sections obtained from mosquitoes fed on mice with or without intravenous rAgApyrase supplementation and used P-selectin (CD62P) staining as a marker for platelet activation. Our results show a significant reduction in P-selectin staining (dark-brown spots) in the midguts of mosquitoes fed on rAgApyrase supplemented mice when compared to the control mice (Fig. 3A and Fig. S11). Quantification of P-selectin staining also showed a significant decrease of 30% in both the mean signal and percentage area of staining for midguts obtained from mosquitoes fed on rAgApyrase supplemented mice (Fig. 3B and 3C). Scanning electron microscopy images showed reduced platelet aggregation in the midguts of mosquitoes fed on rAgApyrase supplemented mice when compared to midguts without apyrase supplementation in which the platelet aggregates were easily distinguished (Fig. 3D and Fig. S8). In the apyrase treated group, platelet aggregates were rarely found and when observed, they were mostly present at the 4 h timepoint (Fig. 3D, and Fig. S8Gi and S8Gii), whereas most of the observed structures resembling platelets at the 30 min timepoint seemed like cells with compromised structure (Fig. 3D, yellow arrows). Interestingly, when mosquitoes were fed on rAgApyrase immunized mice, a hyper-activation and aggregation of platelets was observed, including platelet fragmentation into smaller vesicles and increased filopodia (Fig. 3E, and Fig. S10L, S10M, and S10N) in contrast to the adjuvant immunized group. A similar phenomenon was previously described in an *in vivo* model for thrombus formation where the tightest regions of the thrombus were composed of highly activated and tightly packed platelets that fragmented into vesicular bodies [41].

**Figure 3:**
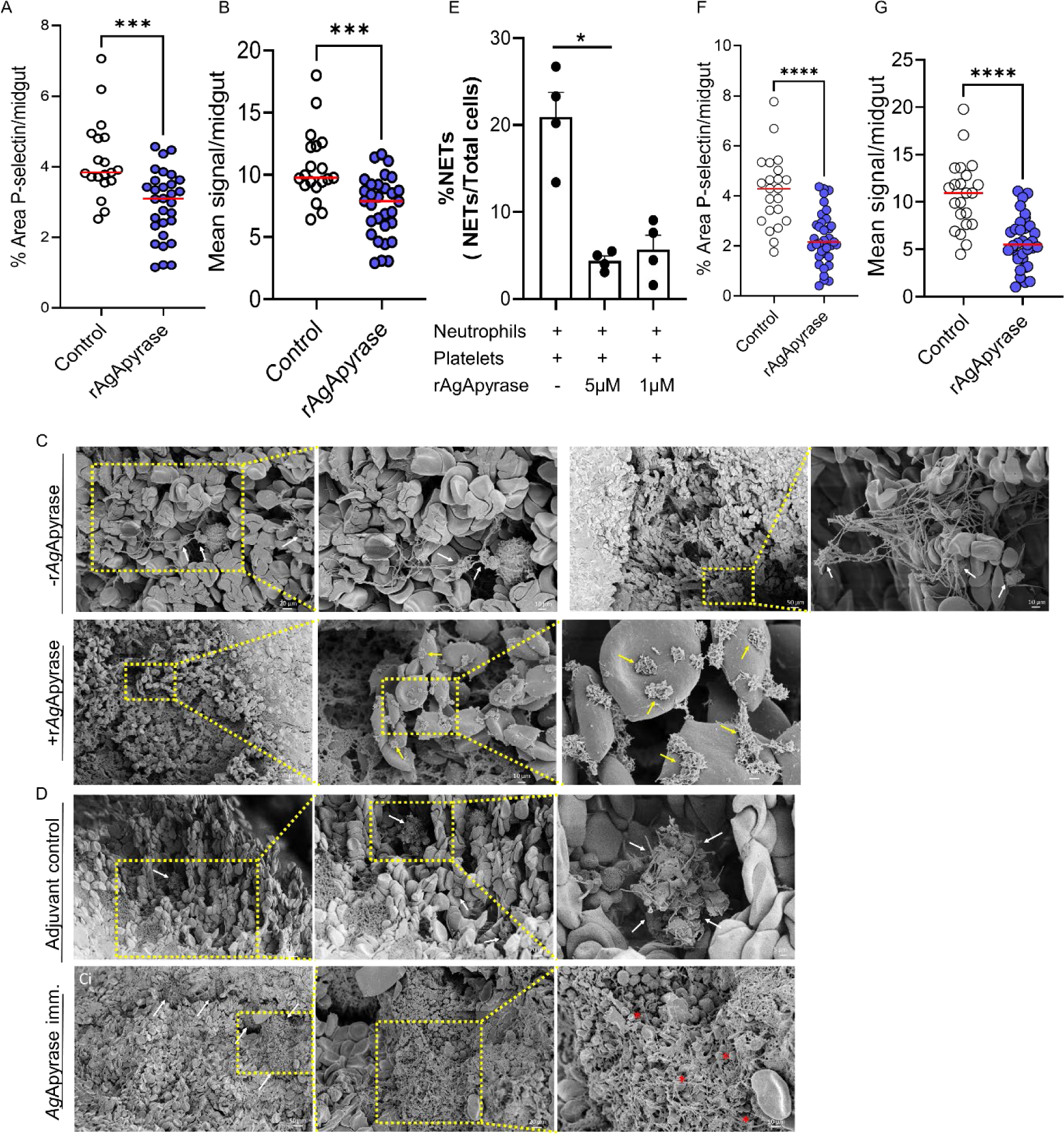
Salivary AgApyrase inhibits platelet activation and aggregation in the blood bolus and decreases *in vitro* NETosis. **(A, B)** Immunohistochemistry was performed on mosquito blood boluses before and after supplementation with rAgApyrase and stained with P-selectin (a marker for platelet activation). Quantification of the percentage area covered by P-selectin staining (A) and the mean P-selectin signal (B) per midgut (from IHC shown in Fig. S11). Data pooled from two independent experiments. Mann Whitney test, ***P=0.0001. **(C, D)** SEM showing platelet aggregation in blood boluses before and after supplementation with rAgApyrase (C) or in from mosquitoes fed on adjuvant or rAgApyrase immunized mice (D). White arrows indicate the aggregation of platelets containing filopodia. Yellow arrows show individual activated platelets that are not aggregated. Red asterisks show fragmentation of platelets into smaller vesicular bodies reminiscence of hypercoagulation. **(E)** rAgApyrase decreases *in vitro* NETosis in the presence of platelets. NETs were quantified by immunofluorescence microscopy with an anti-MPO antibody (Fig. S12A) and the percentage of NETs was calculated as an average of 5-10 fields (400X) normalized to total number of neutrophils. Results expressed as mean % ± SEM. Kruskal-Wallis Test, *P=0.01. (**F, G**) Immunohistochemistry as discussed in panel (A) was performed to quantify neutrophil elastase as a marker for NETosis. From IHC shown in Fig. S13. Data pooled from two independent experiments. Mann Whitney test, ****P=0.0001.

In humans, immunothrombosis involves the cooperative action of the immune and coagulation systems to target invading pathogens. Neutrophil extracellular traps (NETs) play a role in this process by trapping and killing pathogens through the release of decondensed chromatin, histones, myeloperoxidase and elastase. Activated platelets and fibrin induce NET formation (NETosis) and in a positive feedback loop, the negatively charged NETs in turn activates the contact phase of blood coagulation, generating a second positive loop that activates platelets and enhance fibrin polymerization [42, 43]. We investigated the influence AgApyrase on NETosis in the presence of platelets. We found that rAgApyrase, in the presence of platelets, significantly reduces NET formation by 79% (Fig. 3E, Fig. S11). We also observed that rAgApyrase reduces the formation of neutrophil reactive oxygen species (ROS), which are crucial for NETosis (Fig. S12B) [44]. Additionally, immunohistochemistry analysis showed a significant decrease in neutrophil elastase, a marker for NETs, in the mosquito blood bolus after rAgApyrase supplementation (Fig. 2F and 2G, Fig. S13). These findings suggest that salivary apyrase inhibits immunothrombosis in the mosquito midgut, thus protecting the mosquito vector and the malaria parasite.

In this study, we showed that salivary apyrase from *An. gambiae* enhances fibrinolysis and inhibits platelet activation and aggregation in the midgut. We previously demonstrated that fibrin polymerization increases the viscosity of the blood bolus, which in turn reduces parasite motility and transmission to the mosquito [3]. Therefore, we hypothesize that salivary apyrase might enhance parasite transmission to the mosquito by reducing the blood bolus viscosity. To test this hypothesis, we performed passive administration of rAgApyrase on mice infected with *Plasmodium berghei* parasites. A group of *An. gambiae* female mosquitoes were fed on the infected mouse before (control group) and after rAgApyrase intravenous injection. Supplementation of the infected mouse blood with rAgApyrase significantly increased the number of oocysts in the mosquito, while heat-denatured recombinant protein failed to increase the infection (Fig. 4A and S14). Furthermore, immunization of mice with rAgApyrase reduced the oocyst intensity and infection prevalence in mosquitoes when compared to mice immunized with Magic™ Mouse adjuvant alone (Fig. 4B and 4C, and S15). The median oocyst numbers in mosquitoes fed on rAgApyrase-immunized mice ranged between 0 to 4, compared to 3 to 25 in control mice (Fig. 4B and S15). The median infection prevalence was reduced from 81% in control mice to 49% in rAgApyrase-immunized mice (Fig. 4C). Our results show that salivary apyrase ingested by the mosquito during blood feeding increases parasite infectivity to the mosquito.

**Figure 4:**
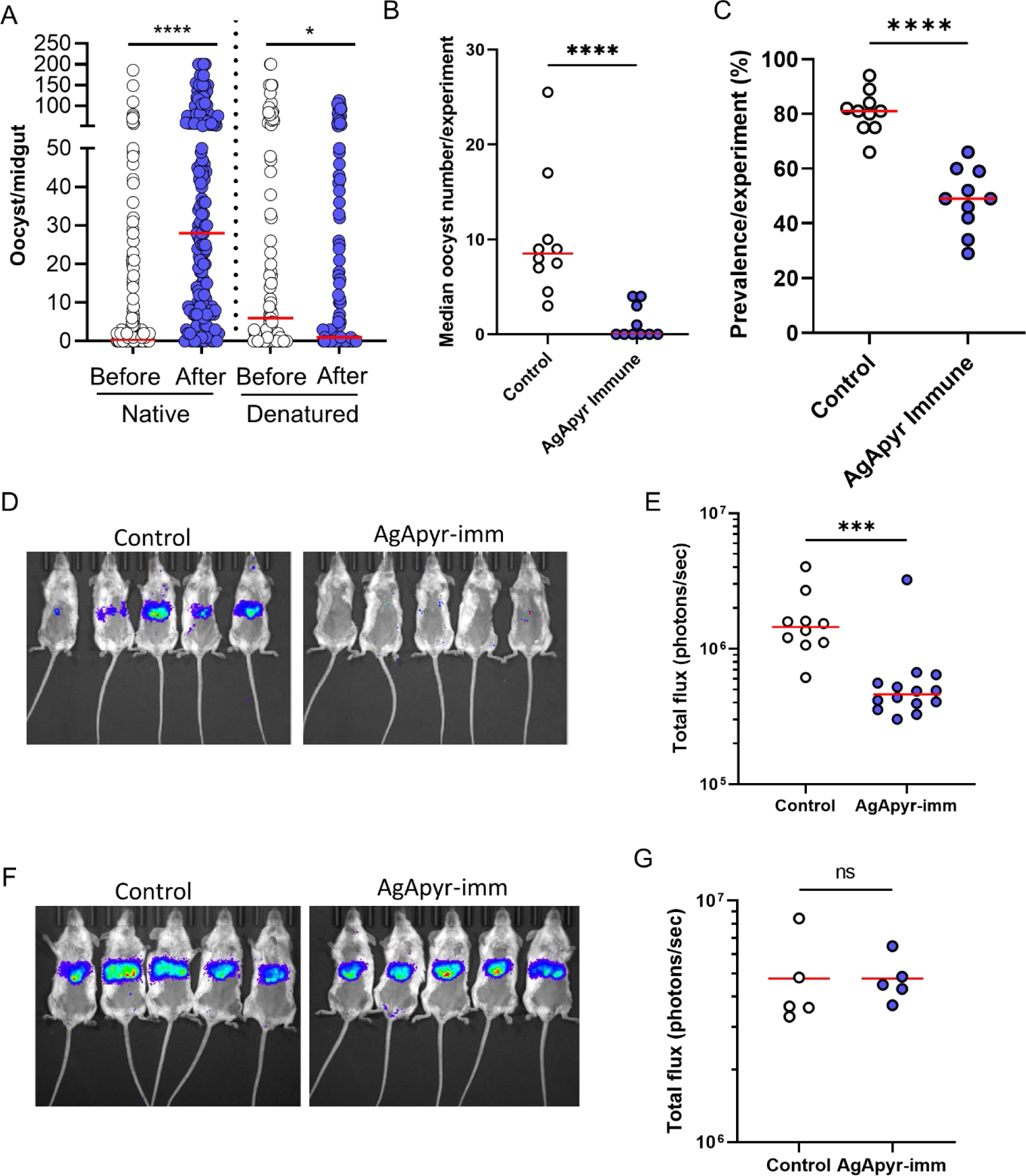
Effect of *Ag*Apyrase on *P. berghei* transmission. **(A)** AgApyrase facilitates *P. berghei* infection of mosquito midguts. Oocyst numbers from midguts of *An. gambiae* mosquitoes that fed on a *P. berghei* infected mouse before or after the intravenous injection of native or heat-denatured rAgApyrase. Data pooled from three individual experiments shown in Fig. S14, and groups were compared with two-tailed t-test followed by Mann-Whitney comparison test. Red lines indicate median. ****P <0.0001; *P<0.05. **(B)** Ag Apyrase immunization inhibits *P. bergei* midgut infection. Oocyst numbers were determined in the midguts of *An. gambiae* mosquitoes fed on *P. berghei* infected BALB/c mice previously immunized with rAgApyrase using Magic Mouse adjuvant or with adjuvant alone as control. Each dot represents median oocyst number from mosquitoes fed on one mouse. Groups were compared with two-tailed t-test followed by Mann-Whitney comparison test. Red lines indicate median. ****P <0.0001. The oocysts numbers from mosquitoes feeding on each individual mouse are shown on Fig. S15. **(C)** AgApyrase immunization reduces prevalence of *Plasmodium* midgut infection. The data obtained from the previous experiment was used to calculate the prevalence of infection. Each dot represents the prevalence of infected mosquitoes fed on one animal. Groups were compared with two-tailed t-test followed by Mann-Whitney comparison test. Red lines indicate median. ****P <0.0001. **(D and E)** AgApyrase immunization inhibits sporozoite transmission. BALB/c mice were immunized with rAgApyrase in Magic Mouse adjuvant or adjuvant alone (control). Mice were challenged with the bite of five *An. stephensi* mosquitoes infected with *P. berghei* sporozoites expressing the luciferase gene. Sporozoite infectivity was determined by measuring luciferase activity in the mouse liver 40 h post challenge. Luminescence signal in the mice livers is shown in panel D and the quantification in panel E. Data pooled from two independent experiments and groups were compared with two-tailed t-test followed by Mann-Whitney comparison test. Red lines indicate median. ***P <0.0002. **(F and G)** Similar experiment as in panels D and E, but mice were challenged with 2000 sporozoites injected intravenously. Data from a single experiment with five mice per group.

*Plasmodium* infection does not reduce the volume of saliva but alters the quality of saliva secreted by a mosquito [45]. Mosquitoes with sporozoites bite more frequently than uninfected mosquitoes and one of the possible ways this occurs is that sporozoites reduce the apyrase activity in the mosquito saliva [46]. The effect of sporozoite infection on mosquito saliva secretion and its impact on apyrase activity has been studied but there are no reports detailing the effect of salivary apyrase on sporozoite transmission. We further studied whether salivary *Ag*Apyrase is required for sporozoite transmission from the mosquito to the mammalian host. Mice immunized with r*Ag*Apyrase were challenged with the bite of five mosquitoes infected with transgenic *P. berghei* sporozoites expressing the reporter luciferase gene. Analysis of the parasite’s luciferase signal in the mouse liver 40 h post-challenge served as an indicator of sporozoite infectivity (escape from the skin and liver invasion) and exoerythrocytic form development [47]. We observed a significant reduction of 68% in luminescence in mice immunized with rAgApyrase compared to adjuvant treated mice (Fig. 4D and 4E). This reduction resulted in a delayed time to patency of 1.5 days (Fig. S16). Notably, when sporozoites were injected intravenously to bypass the skin barrier, the parasite liver load between adjuvant and rAgApyrase immunized mice remained unchanged (Fig. 4F and 4G), suggesting that rAgApyrase immunization affects sporozoite infectivity in the skin. These findings show that salivary AgApyrase is a critical factor for successful sporozoite transmission *via* the mosquito bite.

Our research sheds light on the crucial role of mosquito salivary apyrase in regulating hemostasis and promoting *Plasmodium* development in the mosquito midgut and sporozoite transmission during a mosquito bite. These findings emphasize the role of saliva proteins during malaria parasite transmission and identify salivary apyrase as a unique candidate for a transmission-blocking vaccine, targeting transmission from the vector to the human and from the human to the mosquito vector. Vector saliva regulates hemostatic responses and pathogen transmission at the bite site, however the understanding of how the vector saliva regulates these responses in the mosquito blood bolus and its impact on *Plasmodium* transmission is limited [34].

We previously showed that activation of the coagulation system in the mosquito midgut increases the viscosity of the blood bolus, which serves as a barrier that reduces parasite infectivity to the mosquito [3]. The parasite overcomes this barrier by hijacking plasminogen activators (tPA and uPA) and plasminogen to activate it into plasmin, a protease that degrades fibrin [3]. Likewise, *Plasmodium* sporozoites hijack plasmin to degrade the extracellular matrix of the dermis, facilitating infection of the mammalian host [3]. Here, we show that mosquito salivary apyrase is ingested in high quantities during blood feeding, activates fibrinolysis through the activation of tPA, reduces coagulation by inhibiting platelet activation and aggregation, and inhibits NETosis in the blood bolus. Therefore, salivary apyrase is a key player in preventing hemostasis and inflammation in the blood bolus, properties that will facilitate blood digestion, enhance mosquito fitness and therefore, enable the establishment of parasite infection in the mosquito midgut (Fig. S17). It is possible that apyrase also facilitates sporozoite transmission by enhancing ECM degradation at the dermis through the activation of the fibrinolytic system, although additional experiments are needed to validate this hypothesis. Nonetheless, apyrase has a critical role in facilitating sporozoite transmission to the mammalian host. To the best of our knowledge, this is the first report providing clear mechanistic evidence on the crucial role of mosquito saliva during host-to-vector and vector-to-host transmission of malaria parasites.

Blood feeding by mosquitoes is a fast process compared to other hematophagous vectors, taking only a few minutes. Although *Anopheles* mosquitoes can feed from blood pools, they feed more frequently by cannulating venules or arterioles [48]. The benefits of saliva at the bite site are more apparent during pool feeding compared to direct vessel feeding. Mosquito saliva is primarily assumed to exert its anti-hemostatic and anti-inflammatory effects on the host skin during probing. Interestingly, studies have shown that saliva is expelled throughout the entire blood feeding process in Anopheline mosquitoes, ensuring a constant flow of saliva and salivary proteins [49]. A previous study using *An. campestris* mosquitoes found that the most abundant salivary proteins, including apyrase, were substantially depleted after ingestion of a blood meal, showing that a significant amount of saliva is delivered during blood feeding [50]. Our results show that during blood feeding, a substantial amount of the expelled saliva is ingested and plays a major role in regulating hemostasis in the blood bolus and facilitating *Plasmodium* development in the mosquito.

Our findings underscore the importance of investigating the diverse roles assigned to mosquito saliva, not only during probing and feeding on the host skin but also within the mosquito blood bolus, where it can significantly impact parasite transmission. Studying the functional roles of saliva proteins within this microenvironment can reveal critical insights into the complex interactions between the mosquito, host, and parasite during transmission paving the way for the development of novel strategies for blocking the transmission of not only malaria but also other vector-borne diseases.

## Materials and Methods

### Animal ethics protocol

The study was performed in strict accordance with the recommendations from the Guide for Care and Use of Laboratory Animals of the National Institutes of Health (NIH). The animal use was done in accordance with the Use Committee or The NIH Animal Ethics Proposal SOP LMVR 22.

### Mosquito rearing and *Plasmodium berghei* infection

*Anopheles gambiae* G3 and *An. stephensi* (Nijmegen) mosquitoes were reared at 27°C and 80% humidity with a 12-hour light/dark cycle under standard laboratory conditions and maintained with 10% Karo syrup solution during adult stages. *P. berghei* infections were performed using a transgenic line expressing mcherry and luciferase [51] and maintained by serial passages in female Swiss webster mice. Parasitemia was assessed by light microscopy by using methanol-fixed blood-smears from infected mice and stained with 10% Giemsa. Female mosquitoes (4-5 days old) were fed on infected mice once they reached 3-5% parasitemia. After feeding, infected mosquitoes were kept at 19°C, 80% humidity and 12-hour light-dark cycle. To determine the oocyst numbers, midguts were dissected 10 days post-infection, stained with 0.2% mercurochrome, and mature oocysts were counted by light microscopy.

### Salivary gland dissection and saliva collection

Sugar-fed female adult mosquitoes (4–5 days old) were anesthetized with CO_2_, transferred to an ice-chilled plate, and their salivary glands were dissected under a stereomicroscope in 1X PBS (137 mM NaCl, 2.7 mM KCl, 4.3 mM Na_2_HPO_4_, and 1.4 mM KH_2_PO_4_, pH 7.4). Salivary gland extract (SGE) was obtained by sonicating the glands on a Branson Sonifier 450 sonicator with the probe immersed for 5cm in a 100ml beaker filled with 80ml ice-cold water. The 1.5 ml tube containing the glands was held in this beaker with the help of forceps such that the base of the tube was under the tip of the probe. Power was set at 6 and three rounds of 50% cycle were run for 1 min. Tissue debris were cleared by centrifugation at 10,000 g for 10 min at 4 °C. The supernatants were stored at -80 °C until used [52]. Saliva collection was done by placing the back of live mosquitoes on double-sided tape. Mosquito mouthparts (proboscis) were inserted into 10 μl pipette tips containing 10 μl of 1X PBS. Mosquitoes were allowed to salivate for 20 min at room temperature. The saliva was stored at -80 °C until used.

### Single-chain tPA activity assay

Activation of sc-tPA (100 nM, Innovative Research, Novi, MI, USA, #IHUTPA85SC100UG) was tested in the presence of salivary gland extracts (one, two and four pairs), saliva (10, 20 and 40 μl), or recombinant proteins (100nM) in 0.01% Tween 20, tris-buffered saline (TBST) (50 mM Tris-Cl, pH 7.5 and 150 mM NaCl]. The tPA fluorogenic substrate D-Val-Leu-Lys 7-amido-4-methylcoumarin (10 μM) (Millipore-Sigma, Burlington, MA, USA, #A8171) was added, and the change in fluorescence (excitation, 280; emission, 460) was recorded every 1 min over a period of 4 hours at 37°C using the Cytation 5 microplate reader (BioTek Instruments, USA).

### Salivary gland fractionation and mass-spectrometry

Salivary gland extracts obtained by sonication from 400 pairs of salivary glands were subjected to size-exclusion chromatography on a Superdex 200 10/300 GL column (Cytiva, Marlborough, MA, USA, #28-9909-44) and 1 ml fractions were collected. Each fraction was tested in the sc-tPA activity assay and the fractions showing activity along with five different fractions showing no activity were analyzed by mass spectrometry at the Protein Chemistry Section of the Research and Technology Branch (NIAID, NIH). Proteins in the samples were reduced, alkylated, digested using trypsin, and cleaned up with sold-phase extraction tips (OMIX10, Agilent). The peptide samples were loaded to a PepMap 100 C18 trap column (Thermo Scientific, 75 µm diameter, 2 cm length, 3 µm particles), and separated using a PepMap C18 analytical column (Thermo Scientific, 75 µm diameter, 25 cm length, 2 µm particles) using a 2-hour acetonitrile gradient (0 – 40% in 100 min, 40 – 80% in 5 min, hold at 80% for 5 min, 80 – 0 % in 5 min, and hold at 0% for 5 min) with the flow rate of 300 nL/min using EASY nLC 1000 (Thermo Scientific). The MS data acquisition was carried out in a data-dependent acquisition mode using Orbitrap Fusion™ Lumos™ Mass Spectrometer (Thermo Scientific). The survey MS1 scans were recorded every 2 seconds with the Orbitrap mass analyzer at the 120,000-resolution setting. For the multiple-charged precursors, ddMS2 was performed with the quadrupole isolation of m/z 1.6 window, dissociated by CID, and scanned using the Linear Ion Trap mass analyzer. The dynamic exclusion was utilized for a duration of 30 seconds. The data were analyzed using PEAKS X plus against the *Anopheles gambiae* protein database from VectorBase.org (VB2019-06) and cRAP commonly contaminating protein database from theGPM.org. All peptides were filtered at a 0.5% FDR, and proteins identifications required a minimum of 2 peptides.

### Cloning, expression, and purification of recombinant candidate proteins

DNA sequences of eight candidate proteins were codon-optimized for mammalian expression systems, synthesized, and cloned in the plasmid VR2001-TOPO (Vical Incorporated, San Diego, CA, USA) by BioBasic Inc. The plasmids harboring the desired candidate genes were transformed in One Shot TOP10 Chemically Competent *E. coli* (Invitrogen, Waltham, MA, USA, # C404010). Plasmid DNA was purified using NucleoBond PC 2000 plasmid megaprep kit (Macherey-Nagel, #740549). Recombinant protein expression was carried out at the Protein Expression Laboratory, Advanced Technology Research Facility (NCI, Frederick, MD, USA) Briefly, FreeStyle 293F cells (Thermo Fisher Scientific, Waltham, MA, USA, #R79007) were transfected with 1 mg of plasmid DNA, and supernatants were collected after 72 h of transfection. His-tagged recombinant proteins were purified from the supernatants by affinity chromatography followed by size-exclusion chromatography, using Nickel-charged HiTrap Chelating HP (Cytiva, Marlborough, MA, USA, #17040903) and Superdex 200 10/300 GL columns, respectively (Cytiva, Marlborough, MA, USA, #28-9909-44). All protein purifications were carried out using the AKTA pure system (Cytiva, Marlborough, MA, USA). Purified proteins were resolved in a NuPAGE Novex 4–12% Bis-Tris protein gels (Thermo Fisher Scientific, Waltham, MA, USA, # NP0321BOX) and visualized by Coomassie blue stain using the eStain protein stain system (GenScript, Piscataway, NJ, USA). Protein identity was verified by Edman degradation by the Protein Chemistry Section of the Research Technologies Branch, NIAID, NIH.

### AgApyrase interaction with sc-tPA

To analyze the interaction of AgApyrase with sc-tPA, 96 wells flat-bottom clear plates (Costar, Corning, NY, USA, #COS3915) were coated overnight at 4°C with 20nM rAgApyrase and an unrelated protein amongst the eight candidate proteins (AGAP007393) in carbonate bicarbonate buffer pH 9.6 (Millipore-Sigma, Burlington, MA, USA, #SRE0102). Plates were washed with 0.05% *v*/*v* Tween-20 in TBS (25 mM Tris, 150 mM NaCl, pH 7.4) between all steps before using the stop solution (2N H_2_SO_4_). Plates were blocked with blocking buffer (TBS-Tween with 5% Skim milk) for 1 h at 37°C. After the blocking step, plates were coated with sc-tPA (20nM) and incubated at 37°C for 2 h. Anti-sc-tPA (Innovative Research, Novi, MI, USA, #ISHAHUTPAAP100UG) at 1:1500 dilution was used as a primary antibody to detect protein-protein interactions. Primary antibodies were incubated for 1 h at 37°C. Donkey anti-sheep IgG labeled with horse radish peroxidase (Abcam, Waltham, MA, USA, #ab6900 (1:2500) in blocking buffer was added to the wells and the plates were incubated for 1 h at 37°C. Plates were developed with 1-Step ultra TMB-ELISA substrate solution (Thermo Fisher Scientific, Waltham, MA, USA, #34028) and incubated at room temperature for 15-30 minutes. 2N H_2_SO_4_ solution was added to stop the reaction and absorbance was measured at 450 nm using Cytation 5 microplate reader (BioTek Instruments, USA).

### Plasmin activity assay

Plasmin activity was measured using the specific plasmin chromogenic substrate D-Val-Leu-Lys 4-nitroanilide dihydrochloride (Millipore-Sigma, Burlington, MA, USA, #V0882). All proteins were prepared in phosphate-lysine buffer (10mM potassium phosphate, 70mM sodium phosphate, 100mM lysine, pH 7.5). Phosphate-lysine buffer, rAgApyrase (100nM), sc-tPA (100nM), tc-tPA (100nM), plasminogen (400nM), and rAgApyrase with plasminogen were used as negative controls. Plasmin (400nM), sc-tPA and tc-tPA with plasminogen were used as positive controls. The respective samples were added to a 96-well flat bottom clear plate and 3.58mg/mL of chromogenic substrate were added. Plasmin activity was monitored by continuously measuring the rate of change of absorbance at 405 nm for 20 minutes using Cytation 5 microplate reader (BioTek Instruments, USA).

### AMP hydrolysis activity assay

A malachite green assay kit (R&D Systems, Minneapolis, MN, USA, #DY996) was used to determine the phosphate released by rAg5’NTE using AMP (Millipore-Sigma, Burlington, MA, USA, #A1752) as a substrate. The manufacturer’s microplate assay protocol was followed. Briefly, phosphate standards from 0 μM to 100 μM were prepared and 50 μl of CD73 (0.1 μg) (Millipore-Sigma, Burlington, MA, USA, # N1665) was used as a positive control. 10 μl malachite green reagent A was added, and the plate was incubated for 10 min at room temperature. 10 μl of malachite green reagent B was added and the plate was incubated for 20 minutes at room temperature. The absorbance was measured at 620 nm using Cytation 5 microplate reader.

### ATP and ADP hydrolysis activity assay

Apyrase activity was determined by using a micro-colorimetric method for measuring release of inorganic phosphate from ATP or ADP as previously described [53, 54]. Briefly, a buffer containing 50 mM Tris pH 8.3, 150 mM NaCl and 5 mM CaCl_2_, was mixed with potato apyrase (positive control) (Millipore-Sigma, Burlington, MA, USA, #A6410), buffer alone (negative control) and rAgApyrase in the wells of a 96-well microtiter plate with a final volume of 0.1 ml containing 2 mM of either ATP (Millipore-Sigma, Burlington, MA, USA, #A26209) or ADP (Millipore-Sigma, Burlington, MA, USA, #A2754). This mixture was incubated at 37°C for 10 min. The reaction was stopped by the addition of 28 μl of a stop reaction mixture: 3 μl of the reducing reagent (mixture of 0.2 g 1-amino-2-naphtol-4-sulfonic acid (Millipore-Sigma, Burlington, MA, USA, #08751) with 0.2 g sodium bisulfite (Millipore-Sigma, Burlington, MA, USA, #243973) and 1.2 g sodium sulfite (Millipore-Sigma, Burlington, MA, USA, #239321) diluted to give a final concentration of 25 mg/ml) and 25 μl of 1.25% ammonium molybdate (Millipore-Sigma, Burlington, MA, USA, # 277908) in 2.5 N H_2_SO_4_ [53]. The absorbance was measured at 650nm using Cytation 5 microplate reader. The concentration of phosphate was determined by interpolation with a phosphate standard curve.

### Western blot assay

Western blot assay was used (i) to study the interaction of sc-tPA and AgApyrase and (ii) determine the specificity of the polyclonal anti-apyrase antibodies. To study the interaction of sc-tPA with AgApyrase, sc-tPA at different concentrations (200nM, 500nM and 1µM) was incubated with 500nM rAgApyrase at 37 °C for 4 h. sc-tPA and tc-tPA were used as positive controls. To determine the specificity of the poly-clonal anti-rAgApyrase antibodies, human serum (2µl) and human plasma (2µl) along with human blood-fed mosquitoes and unfed mosquitoes were tested with anti-apyrase antibodies. The mosquito midguts were dissected 30 min post blood feeding in PBS supplemented with 5 mM EDTA, 0.1 mM phenylmethylsulfonyl fluoride (Millipore-Sigma, Marlborough, MA, USA), and protease inhibitors (Millipore-Sigma, Marlborough, MA, USA). A total of five midguts were pooled in 50 μl of PBS, then the midguts were macerated on ice. For both the experiments, reducing SDS–polyacrylamide gel electrophoresis (PAGE) sample buffer (LI-COR Biotechnology, Lincoln, NB, USA) was added to the samples and were separated by SDS-PAGE and transferred to polyvinylidene difluoride membranes (Millipore--Sigma, Marlborough, MA, USA). The membrane was blocked with Intercept^®^ (PBS) Blocking Buffer (LI-COR Biotechnology, Lincoln, NB, USA) overnight at 4 °C following incubation with mouse anti-rAgApyrase antibody (1:10,000 in blocking buffer) or rabbit anti-tPA antibodies (1:2500) (Millipore-Sigma, Marlborough, MA, USA). Membranes were incubated with anti-mouse or anti-rabbit horseradish peroxidase (1:2500 in blocking buffer) for 1 hour at room temperature and detected using SuperSignal Western Dura PLUS Chemiluminescent Substrate (Thermo Fisher Scientific).

### Multiple sequence alignment, phylogenetic analysis, and structure prediction

Multiple sequence alignment was done for AgApyrase (gene ID: AGAP011026) with the 5’ NTE sequences available from different mosquito species and hematophagous arthropods using CLUSTAL Omega [55]. To understand the evolutionary position of Ag5’NTE, a phylogenetic tree was constructed using MEGA 6.0 [56]. The evolutionary history was inferred using the maximum-likelihood method [57]. The bootstrap consensus tree inferred from 1000 replicates was taken to represent the evolutionary history of the taxa analyzed [58] where the analysis involved 18 amino acid sequences. All positions containing gaps and missing data were eliminated. To obtain the phylogenetic tree, neighbor-join and BioNJ algorithms were applied to a matrix of pairwise distances estimated using a JTT model, and then the topology with superior log likelihood value was selected [56].

Models of AG-Apyrase-tPA complex were generated using RoseTTAFold CC [59], OmegaFold [60], and AlphaFold2 Multimer [61] using default parameters. Only the AlphaFold2 models resulted in compact arrangements with OmegaFold suggesting some contacts between the two molecules. The models were subjected to further optimization using Yasara [62] version 22.9.24 and the hm_build.mcr homology modeling macro using the AlphaFold and OmegaFold models as templates and default parameters. This procedure improved the model’s overall quality resulting in a final arrangement with a Z-score of -0.914 with both the Apyrase and tPA models in good agreement with known structures. Several sequences of *Anopheles* Apyrases were aligned using Clustal Omega [55] as implemented in JalView version 2.11.2.6 [63].

### *Ex vivo* platelet-aggregation assay

Platelet aggregation assay was performed as described by Martin-Martin et al., 2020 [64]. Platelet rich plasma was obtained from normal healthy donors enrolled in a protocol approved by the National Institutes of Health Clinical Center Institutional Review Board (NIH protocol 99-CC-0168 “Collection and Distribution of Blood Components from Healthy Donors for In Vitro Use”). Blood donors provided written informed consent, and platelets were de-identified prior to distribution. Platelet aggregation was measured using a light transmission aggregometer (Chrono-Log Corporation, Havertown, PA, USA). Briefly, 300 μL of platelet rich plasma, diluted to approximately 2.5 × 10^5^ platelets/μL in HEPES-Tyrode’s buffer (137 mM NaCl, 27 mM KCl, 12 mM NaHCO_3_, 0.34 mM sodium phosphate monobasic, 1 mM MgCl_2_, 2.9 mM KCl, 5 mM HEPES, 5 mM glucose, 1% BSA, 0.03 mM EDTA, pH 7.4) were pre-stirred in the aggregometer for 1 min to monitor pre-aggregation effects. Recombinant AgApyrase (3μM) or Tyrode’s buffer (negative control) were added to the platelet rich plasma and were placed in a Chrono-Log aggregometer model 700 (Chrono-Log Corporation) and stirred at 1200 rpm at 37 °C for 1 min prior to the addition of ADP (0.3μM) as an agonist.

### Anti-AgApyrase polyclonal antibody generation

BALB/c mice were inoculated with 50 μg of recombinant AgApyrase protein in Magic Mouse adjuvant (Creative Diagnostics, Shirley, MA, USA, #CDN-A001). As control, were inoculated with adjuvant only. Mice were boosted three weeks after the primary injection with the same amount of protein. Terminal bleeds were performed at 35 days post first immunization. All injections and terminal bleeds were carried out by the NIAID/NIH animal facility personnel. The antibody levels were determined by coating recombinant AgApyrase (1μg/ml) on a 96-well, flat bottom clear plate (Costar, Corning, NY, USA) overnight at 4 °C. Wells were washed three times in TBS-Tween and blocked with blocking buffer (5% skim milk in 1XTBST). After 1 h, wells were washed three times in TBS-Tween incubated with adjuvant control or AgApyrase serum samples (1:25,000 dilution) in blocking buffer for 1 h. Wells were washed three times and incubated with HRP-labeled anti-mouse IgG H+L (1:2500 dilution in blocking buffer) (Cell Signaling Technology, Danvers, MA, USA, #7076P2). Plates were developed with 1-Step ultra TMB-ELISA substrate solution and incubated at room temperature for 15-30 min. 2N H_2_SO_4_ solution was added to stop the reaction and absorbance was measured at 450 nm using Cytation 5 microplate reader.

### Immunohistochemistry of mosquito midguts

Immunohistochemistry was performed on either whole mosquitoes or dissected midguts to determine the ingestion of salivary apyrase by the mosquito. Unfed whole mosquitoes (2 days old, negative control) and blood-fed mosquito midguts (human blood NIH protocol 99-CC-0168 “Collection and Distribution of Blood Components from Healthy Donors for In Vitro Use”) were dissected 30 min post-feeding. The mosquitoes and mosquito midguts were fixed in 10% neutral buffered formalin, processed with Leica ASP6025 tissue processor (Leica Microsystems), embedded in paraffin and sectioned at 5 μm for histological analysis. These formalin-fixed paraffin-embedded tissues sections were used to perform immunohistochemical staining using mouse anti-apyrase antibody (1:10,000) to stain unfed and blood-fed mosquitoes. Antibody diluent (Agilent, Santa Clara, CA, USA, # S302283-2) was used as a negative reagent control to replace the primary antibodies. Staining was carried out on the Bond RX (Leica Biosystems) platform according to manufacturer-supplied protocols. Briefly, 5µm-thick sections were deparaffinized and rehydrated. Heat-induced epitope retrieval (HIER) was performed using Epitope Retrieval Solution 1, pH 6.0, heated to 100 °C for 20 min. The specimen was then incubated with hydrogen peroxide to quench endogenous peroxidase activity prior to applying the primary antibody. Detection with DAB chromogen was completed using the Bond Polymer Refine Detection kit (Leica Biosystems, #DS9800). Slides were finally cleared through gradient alcohol and xylene washes prior to mounting and placing cover slips. Sections were examined by a boarded-certified veterinary pathologist using an Olympus BX51 light microscope and photomicrographs were taken using an Olympus DP73 camera.

To determine platelet aggregation inside the mosquito midgut, a group of mosquitoes (labeled as “Before” in the figures) were fed on Swiss-webster mice for 15 min. The same mouse was IV injected with 200 μg of rAgApyrase and a second group of mosquitoes (labeled as “After” in the figures) were fed on it for 15 min. Mosquito midguts were dissected 30 min post-feeding and fixed in 10% neutral buffered formalin and were processed as described above. A rabbit polyclonal CD62p (Bioss, Woburn, MA, USA, #Bs-0561R), which is a marker for platelet activation and aggregation was used to stain the formalin-fixed paraffin-embedded tissues sections with dilution of 1:200. The sections were then processed for detection as described above.

### Scanning electron microscopy

To determine the effect of AgApyrase on fibrin polymerization in the mosquito midgut, a group of mosquitoes (labeled as “Before” in the figures) were fed on Swiss-webster mice for 15 min. The same mouse was IV injected with 200 μg of rAgApyrase and a second group of mosquitoes (labeled as “After” in the figures) were fed on it for 15 min. Mosquito midguts were dissected 30 min and 4 h post-feeding. Mosquito midguts were fixed in 2.5% glutaraldehyde / 4% paraformaldehyde in 0.1M sodium cacodylate buffer. The samples were rinsed in 0.1 M sodium cacodylate buffer three times for 5 min each and postfixed with 0.5% OsO_4_ / 0.8% K_4_Fe(CN)_6_ in 0.1 M cacodylate buffer for 1 h. Following three more 5-min rinses in 0.1 M cacodylate buffer the midguts were dehydrated in a graded ethanol series. The midguts were dropped in liquid nitrogen and cut in half with a razorblade. The midgut halves were placed back into 100% ethanol and dried in a Bal-tec CPD030 critical point drier (Balzers). The samples were mounted on aluminum SEM stubs using carbon sticky tape (Electron Microscopy Sciences, Hatfield, PA, USA) and coated with 12 nm of iridium in an EMS300T D sputter coater (Electron Microscopy Sciences) before being viewed at 2.0 kV in a Hitachi SU8000 field emission scanning electron microscope (Hitachi, High-Tech in America) in secondary electron mode. The same protocol was also used for midguts from mosquitoes fed on adjuvant control and rAgApyrase immunized mice. Midguts from mosquitoes feeding on immunized mice were dissected 30 min post-feeding.

### Transmission electron microscopy

*An. gambiae* mosquito midguts obtained from “Before” and “After” groups discussed above were fixed in 4% paraformaldehyde in 0.1M phosphate buffer. Midguts were then rinsed in 0.1M cacodylate buffer 3 times for 5 min at room temperature each before being dehydrated in a graded ethanol series from 50% to 90% ethanol. The samples were then infiltrated in a 1:1 mixture of 90% ethanol and LR White resin with accelerator (Electron Microscopy Sciences, Hatfield, PA, USA) for 16 h at 4 °C on a shaker. After infiltrating the tissue in 100% LR White resin for 8 h, the midguts were embedded in fresh LR White resin in gelatin capsules (Electron Microscopy Sciences) at 50 °C under vacuum for 48 h. Thin sections were cut on a Leica UC6 ultramicrotome (Leica Microsystems, Wetzlar Germany) and collected on 200 mesh formvar/carbon coated nickel grids (Electron Microscopy Sciences). Immunolabeling was done using the drop flotation method where grids are floated on 10 to 50 μL droplets of the solution for each step. Grids were blocked with 1% BSA / 1% gelatin / 0.01% Tween20 in PBS for 10 min then labeled with primary antibody (anti-fibrinogen Millipore--Sigma, Marlborough, MA, USA #341552 or anti-TER-119, BD Biosciences, San Jose, CA, USA #550565) diluted 1:25 in blocking buffer for 1 h. The grids were washed twice with blocking buffer before labeling with secondary antibodies (goat anti-rat or goat anti-rabbit, 10nm, Electron Microscopy Sciences), rinsed three times in 1X PBS, and rinsed three additional times in filtered dH_2_O. The grids were stained with 1% uranyl acetate for 8 min and imaged in a Hitachi 7800 transmission electron microscope at 80 kV (Hitachi High-Tech in America).

### D-Dimer assay

To measure D-dimer formation in mosquito blood boluses, mosquito midguts from “Before” and “After” groups mentioned above were used. ELISA (From LifeSpan BioSciences, Mouse Fibrin Degradation Product D-dimer ELISA Kit (LS Bio, #LS-F6179) was used to test the level of D-dimer in 5 and 10 mosquito midguts. The mosquito midguts were dissected in 1X PBS 30 min post feeding, homogenized thoroughly, and subjected to three cycles of freeze (-20 °C)/ thaw (room temperature). The cell debris were removed by centrifugation at 5000g for 5 min at 4 °C and the supernatant was used for the assay. The assay was performed as per manufacturer’s protocol. Briefly, standard solutions from a range of 0, 617.3 to 50000 pg/ml were prepared for a standard curve to which 50 μl of detection reagent A were added and the plate was incubated for 1 h at 37 °C, followed by three washes with the wash buffer. 100 μl of detection reagent B were added to the plate and incubated for 30 min at 37 °C. The plate was washed five times with the wash buffer, 90 μl of TMB substrate solution was added to the plate and incubated for 15 min at 37 °C. The reaction was stopped by adding 50 μl of stop solution and the absorbance was recorded at 450nm using Cytation 5 microplate reader (BioTek Instruments, USA).

### *In vitro* NETosis assay

Peripheral blood was collected in heparinized tubes after venipuncture. Blood was fractionated via density gradient centrifugation using Ficoll-Paque Plus (GE Healthcare). Neutrophils were isolated by dextran sedimentation, washed with PBS 1X, and red blood cells were lysed with hypotonic salt solution. Neutrophils were resuspended in RPMI serum-free at a density of 1×10^6^ cells/mL. For isolation of platelet rich plasma, peripheral blood was collected in heparinized tubes, blood was centrifuge at 1000 rpm for 10 min, then supernatant plasma containing platelets was transfer to a new sterile tube and centrifuged at 3000 rpm for 10 min at room temperature. After centrifugation, 2/3 parts of volume plasma were discarded, and platelets were suspended in a minimum quantity of plasma (2-4 mL) and counted. Neutrophils were preincubated with 1-5 uM of the mosquito rAgApyrase. After pre-incubation, neutrophils were incubated with 2 x10^5^ platelets for 2-3 hours at 37C. Cells were fixed with 4% PFA and stored at 4°C until immunofluorescence analysis was performed.

Fixed neutrophils were washed with PBS, and blocked with 0.2% porcine gelatin (Sigma) for 1h at room temperature. Cells were incubated with primary Ab anti-MPO (Dako); for 30 min in a humid chamber at 37°C. Coverslips were washed three times with PBS and incubated with secondary antibody (Goat anti-rabbit IgG, Invitrogen) for 30 minutes at 37°C. Nuclei were counterstained with Hoechst for 10 min at RT. After three washes, coverslips were mounted on glass slides using Prolong Gold solution (Invitrogen). Slides were visualized in a Zoe microscope. Three pictures of each experiment were taken, and NETs were counted.

### Plasmodium berghei infection

To determine the effect of AgApyrase on *Plasmodium* infection, *P. berghei* infections were performed using a transgenic line expressing mcherry and luciferase [51] and maintained by serial passages in female Swiss webster mice. Parasitemia was assessed by light microscopy by using methanol-fixed blood-smears from infected mice and stained with 10% Giemsa. A group of mosquitoes (Before) were fed on infected Swiss-webster mice for 15 min. The same mouse was IV injected with rAgApyrase (200 μg) and a second group of mosquitoes (After) were fed on it for 15 minutes. This assured that the blood-meal in both the groups had similar parasitemia. For heat-denatured protein assay, the protein was incubated at 65 °C for 30 min and then IV injected. Infected mosquitoes were kept at 19 °C, 80% humidity and 12h light-dark cycle. To determine oocyst numbers, midguts were dissected 10 days post-infection, stained with 0.2% mercurochrome, and mature oocysts were counted with by light microscopy. The same protocol was followed for mosquitoes fed on adjuvant control and rAgApyrase immunized mice.

### Sporozoite transmission assay

For mosquito challenge: Adjuvant control and *Ag*Apyrase immunized mice were challenged with the bite of five *An. stephensi* mosquitoes infected with *P. berghei* sporozoites expressing mCherry and luciferase. The infection status of mosquito salivary glands was verified before the challenge by the presence of strong red fluorescence in the thoracic cavity.

For intravenous sporozoite challenge: Adjuvant control and rAgApyrase immunized mice were anesthetized by peritoneal injection of ketamine (50 mg/kg body weight) and xylazine-hydrochloride (10 mg/kg body weight), and then 5,000 *P. berghei* sporozoites expressing mCherry and luciferase, and isolated from salivary glands, were injected intravenously in the lateral tail veins in a total volume of 200 μl of PBS.

Parasite liver load was determined by measuring luciferase activity in the mouse liver as previously reported [47]. Forty-two h after sporozoite injection or mosquito-bite challenge, the mouse abdominal hair was removed using Nair cream, the challenged mice were intraperitoneally injected with 100 μL of D-luciferin (30 mg/mL) (Perkin Elmer, Waltham, MA, USA, #122799), and then anesthetized in an isoflurane chamber. After mice were immobilized, they were placed in an IVIS Spectrum imager to evaluate bioluminescence by measuring the radiance from the abdomen for 5 min. Mice were returned to their cage to monitor parasite development in the blood by the pre-patency assay. Infection patency was monitored by Giemsa-stained blood smears for 12 days after infection. A single experiment included five mice for each treatment.

## Acknowledgments

The authors are grateful to Andre Laughinghouse and Kevin Lee for insectary support; Allison Booth and DeAndre Bacon for technical support with animals; L. Renee Olano, Glenn Nardone and for support with mass spectrometry analysis.

## Funding

This work was funded by NIH Distinguished Scholars Program and the Intramural Research Program of the Division of Intramural Research (AI001250-01), National Institute of Allergy and Infectious Diseases (NIAID), NIH to J.V.R.; Malaria Research Program Fellowship to Z.R.P.;

## Author contributions

Conceptualization: Z.R.P., T.L.A.S., J.V.R. Methodology: Z.R.P., T.L.A.S., J.V.R.; Investigation: Z.R.P., M.M., B.C., E.P.M., C.C.R., P.C.L-V., I.M-M., Y.F-G., R.E.C., N.S., N.S., I.N.M., D.A.A., D.N.G., E.F., M.J.K., E.C., J.V.R. ; Formal analysis: Z.R.P., and J.V.R.; Writing – original draft: Z.R.P. and J.V.R.; Supervision: J.V.R.;

## Competing interests

All data are available in the manuscript or the supplementary materials.

## Notes

### Competing Interest Statement

The authors have declared no competing interest.

