## Supplementary Material for "*Anopheles* salivary apyrase regulates blood meal hemostasis and drives malaria parasite transmission"

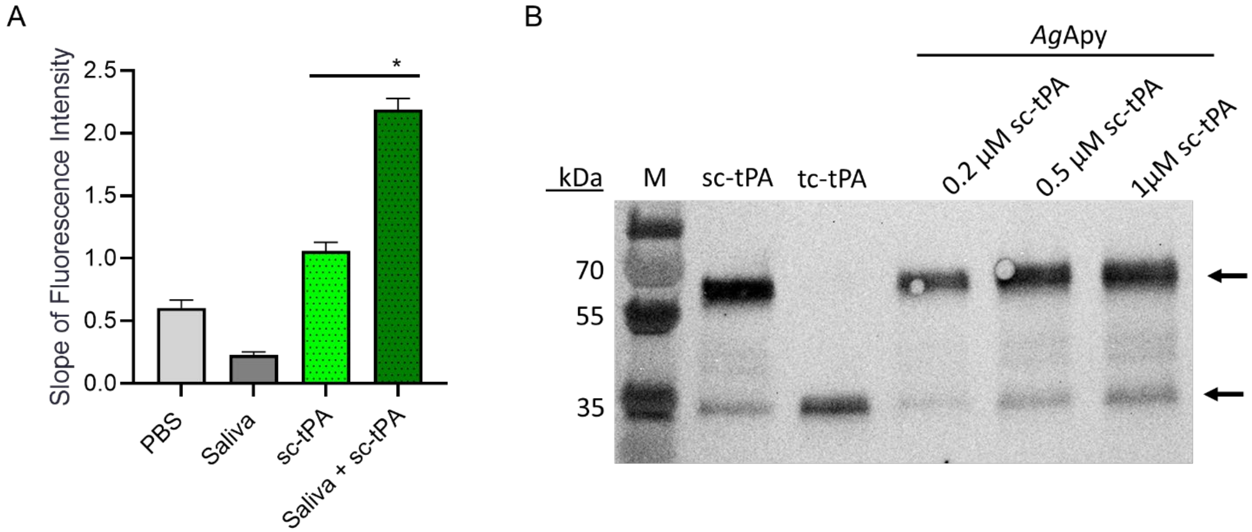


**Fig S1: (A)** Fluorogenic assay for single-chain tPA (sc-tPA) activation showing that mosquito saliva activates sc-tPA. *P <0.01. (**B**) Western blot analysis of sc-tPA after incubation with rAgApyrase. The top arrow indicates sc-tPA (~65kDa) and the bottom arrow indicates tc-tPA (~32kDa) positive controls. sc-tPA at different concentrations (200nM, 500nM and 1μM) was incubated with rAgApyrase to determine the mode of action for tPA activation by AgApyrase. 500 nM of sc-tPA alone were used in the second lane. No extra cleaved bands or an increase in sc-tPA cleavage are seen in any of the sc-tPA samples incubated with rAgApyrase. M: molecular marker.

**A**

| Gene ID | VectorBase annotation | LabID |
|---|---|---|
| AGAP011026 | 5' nucleotidase ecto | ZPJVR1 |
| AGAP011971 | Ser/Thr protein phosphatase/nucleotidase | ZPJVR2 |
| AGAP007393 | protein disulfide isomerase family A member 3 | ZPJVR3 |
| AGAP000610 | SG1f: salivary gland protein 1-like 6 | ZPJVR4 |
| AGAP000376 | Tsf1: Transferrin | ZPJVR5 |
| AGAP012407 | protein disulfide-isomerase A1 | ZPJVR6 |
| AGAP000607 | SG1c: salivary gland protein 1-like 3 | ZPJVR7 |
| AGAP001919 | protein disulfide-isomerase A6 | ZPJVR8 |

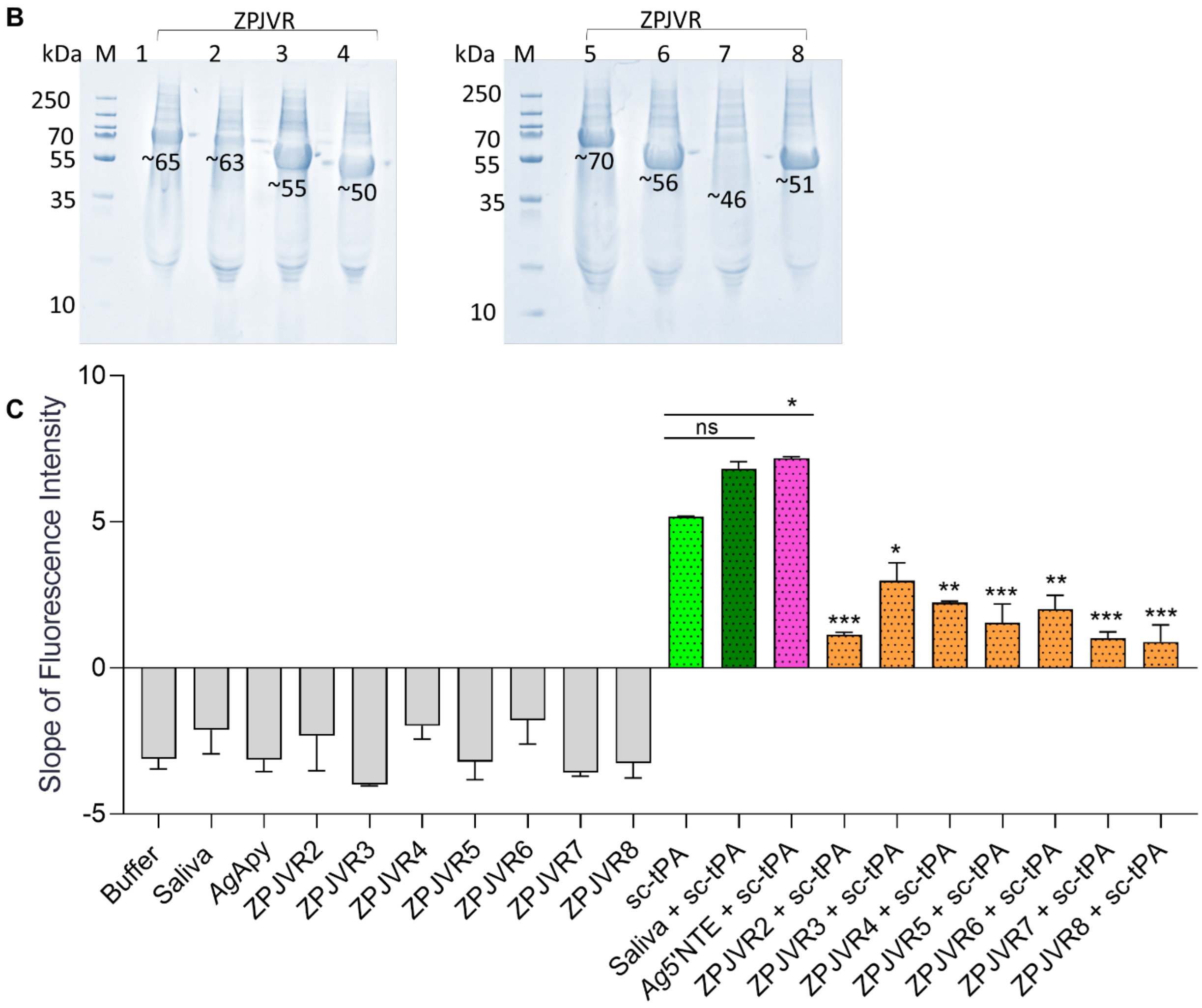


**Fig S2: (A)** Mosquito salivary proteins identified by mass spectrometry of fractions obtained by size-exclusion chromatography were shortlisted as potential tPA activators based on three criteria: 1) present in fractions activating tPA (Z7, Z8, and Z9), 2) must be a secreted protein, and 3) must be absent from fractions that do not activate tPA (A5 and B3). Eight candidate proteins were selected as potential tPA activators (shown in the table). **(B)** Coomassie stained gel of recombinant proteins of the eight tPA activator candidates. ZPJVR 1-8: lab ID for eight candidate proteins; Western blot for AgApyrase using anti-His antibody. M: molecular marker. **(C)** tPA activation fluorogenic assay for the eight identified proteins. All the eight candidate proteins (mustard yellow bars) were tested for tPA activation and only rAgApyrase (pink bar) activated tPA. Data from two independent experiments. Groups were compared with an ordinary one-way ANOVA followed by paired two-tailed t-test. ***P <0.0001; **P <0.001; *P<0.02; ns: not significant.

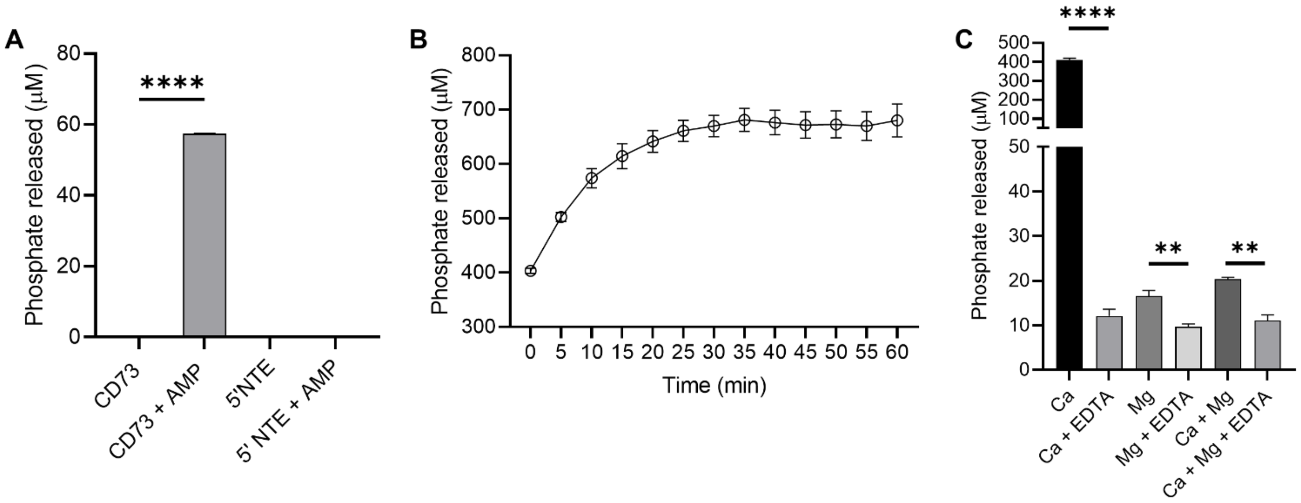


**Fig S3: (A)** Ag5'NTE functions as an apyrase. 5' nucleotidases release inorganic phosphate from AMP. Inorganic phosphate released was measured using malachite green assay. rAg5'NTE did not release phosphate from AMP when compared to the positive control CD73 [human 5'NTE]. Data shown from two independent experiments performed in duplicates. Unpaired t-test, ****P <0.0001. **(B)** Time course of ADP hydrolysis by AgApyrase. Inorganic phosphate released from ADP was measured using malachite green assay over a time-course of 60 minutes. Data shown from one representative experiment performed in duplicates. **(C)** Apyrase activity of rAg5'NTE was measured using malachite green assay in presence of either 5 mM $CaCl_2$ and/or 5 mM $MgCl_2$ with or without addition of 5 mM EDTA. Data shown from a representative experiment performed in triplicates. Unpaired t-test, ****P <0.0001, **P <0.0075.

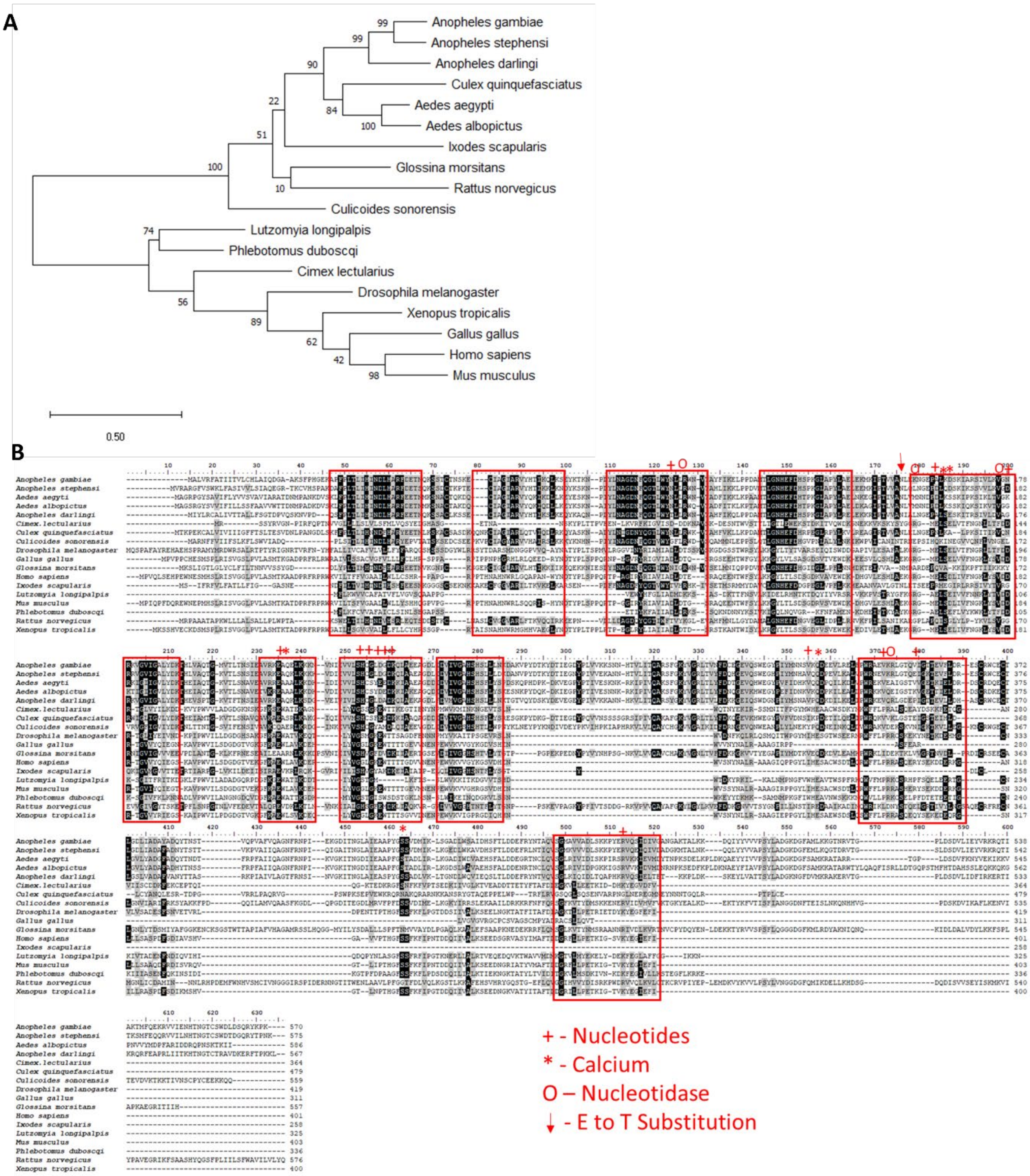


**Fig S4:** **(A)** Phylogeny of apyrase protein sequences. Boot strapping done for 1000 iteration and percentage of trees in which the associated taxa clustered together shown next to the branches. **(B)** Multiple sequence alignment of apyrase sequences. The red boxes depict the conserved region amongst all the species. '+' highlights the residues where nucleotides bind; '*' points the calcium binding site; O points the nucleotidase binding site; ↓ depicts the E to T amino acid substitution in AgApyrase. *Anopheles stephensi* (ASTE008450); *Anopheles darlingi* (ADAC007226); *Culex quinquefasciatus* (CPIJ019168) *Aedes aegypti* (AAEL006347); *Aedes*

*albopictus* (AALF004988); *Ixodes scapularis* (ISC1N003760); *Glossina morsitans* (GMOY012313); *Rattus norvegicus* (AAH81806); *Culicoides sonorensis* (CSON007783); *Lutzomyia longipalpis* (AAD33513); *Phlebotomus duboscqi* (ABI20147); *Cimex lectularius* (AAD09177); *Drosophila melanogaster* (CAL26011); *Xenopus tropicalis* (NP988940); *Gallus gallus* (NP001026752); *Homo sapiens* (NP620148); *Mus musculus* (NP083778).

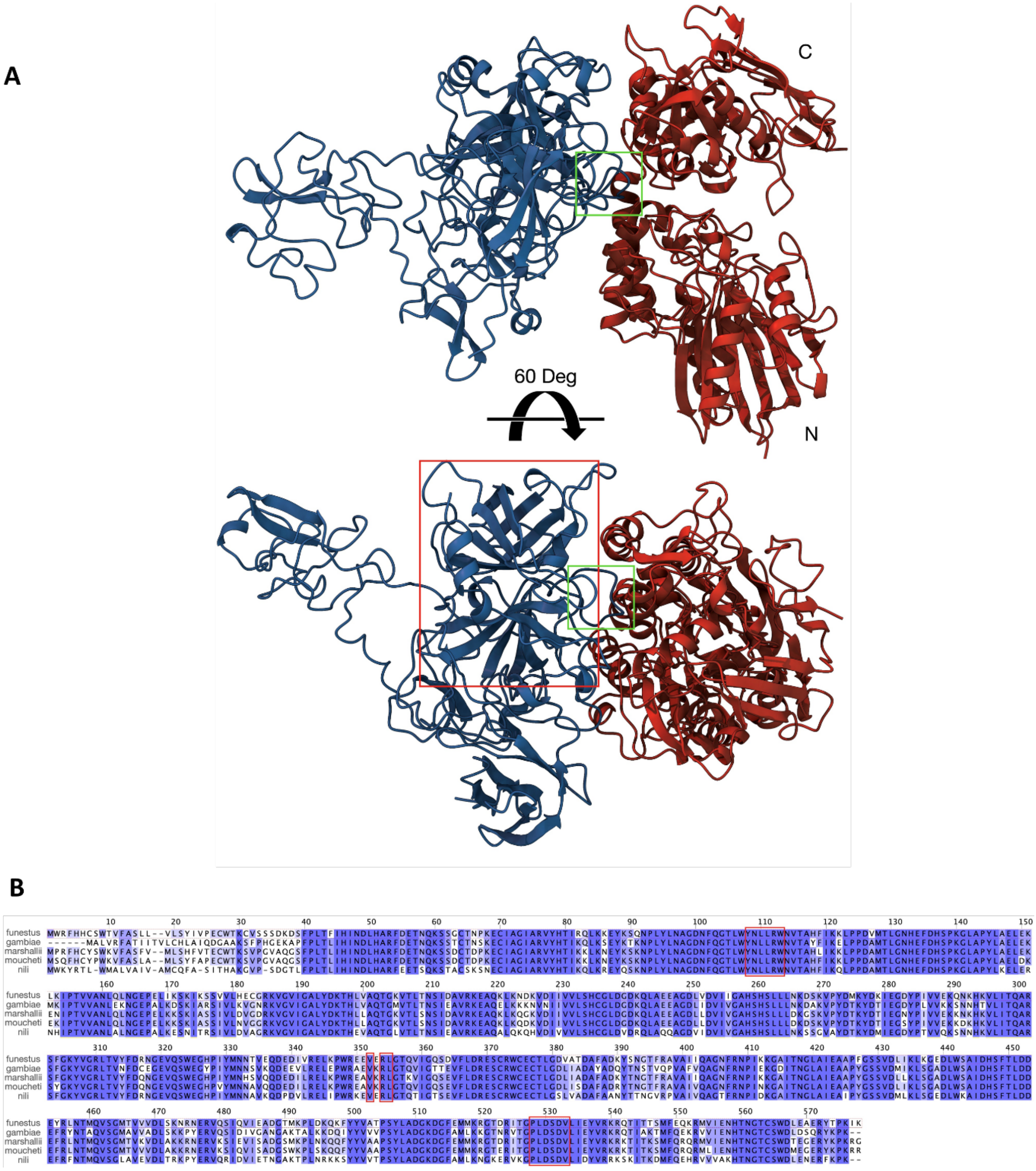


**Figure S5:** Model of the predicted interaction of AgApyrase with human tPA. **(A)** General arrangement of the complex in two orientations, with Apyrase in red (with N and C lobes labeled) and tPA in blue. The buried surface area for the Apyrase-tPA complex final model is 1467 A2. The interaction between the two molecules is mediated by one of tPA the protruding loops (1006-HEALSP-1011, indicated with a green box) in the tPA serine protease domain, which is recognizable and marked by a red box. **(B)** Sequence alignment of apyrases from *An. gambiae*, *An. funestus*, *An. marshallii*, *An. moucheti*, and *An. nili* showing the apyrase residues (highlighted in red boxes) mediating the contact with human tPA. The sequence color indicates the degree of identity.

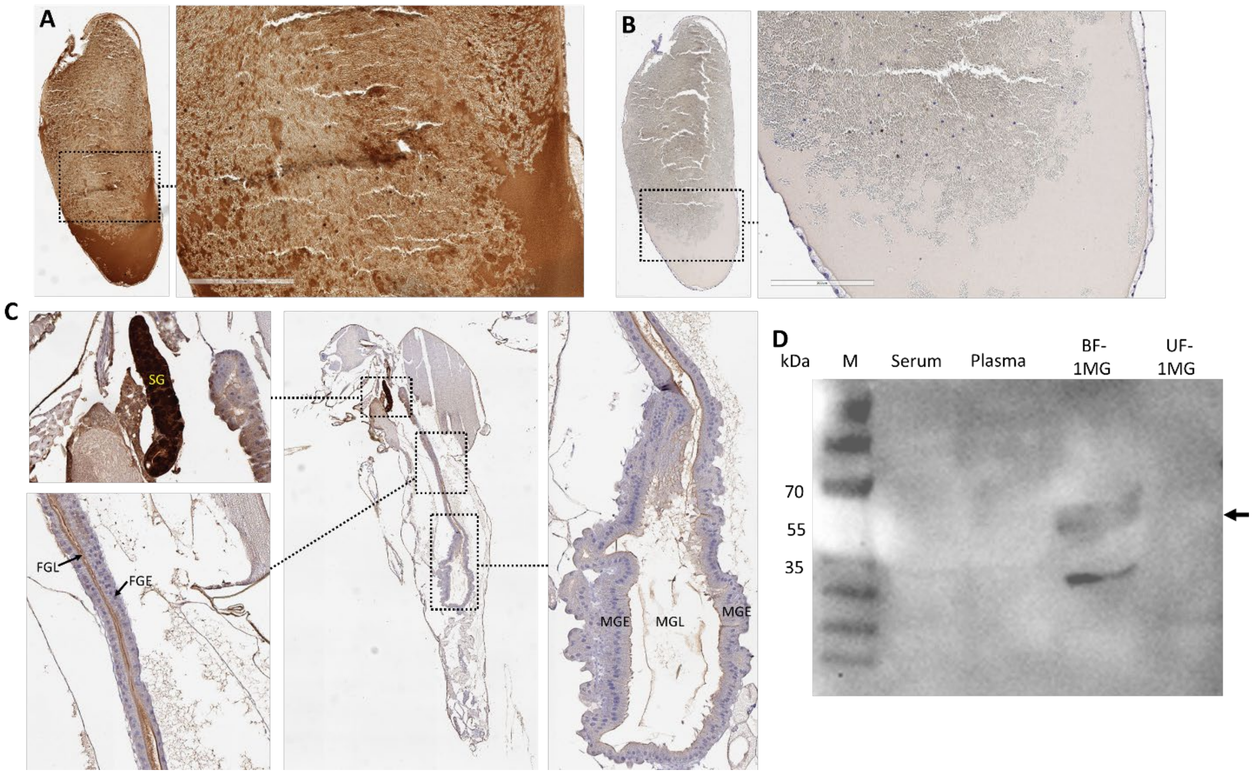


**Fig S6: (A, B)** Immunohistochemistry performed on human blood fed mosquito midgut shows ingestion of salivary AgApyrase which is mixed with the blood bolus content (A, dark brown staining). Panel B shows the staining performed on a different section of the same blood fed mosquito midgut shown in panel A and stained only with secondary antibody. **(C)** Immunohistochemistry performed on unfed whole mosquito with anti-apyrase antibodies showed faint brown staining in the foregut lumen and midgut lumen and strong brown staining in the salivary gland (SG). MGE: midgut epithelium; MGL: midgut lumen; FGL: foregut lumen; FGE: foregut epithelium. **(D)** Western blot analysis showing the specificity of anti-AgApyrase antibodies. The human serum and plasma were tested for the presence of apyrase. No signal was obtained with anti-apyrase antibodies for human serum and plasma and for unfed mosquito midgut. Desired band ~65kDa (arrow) was obtained in blood-fed midgut sample. M: Marker, BF: Blood fed; UF: Unfed; MG: Midgut.

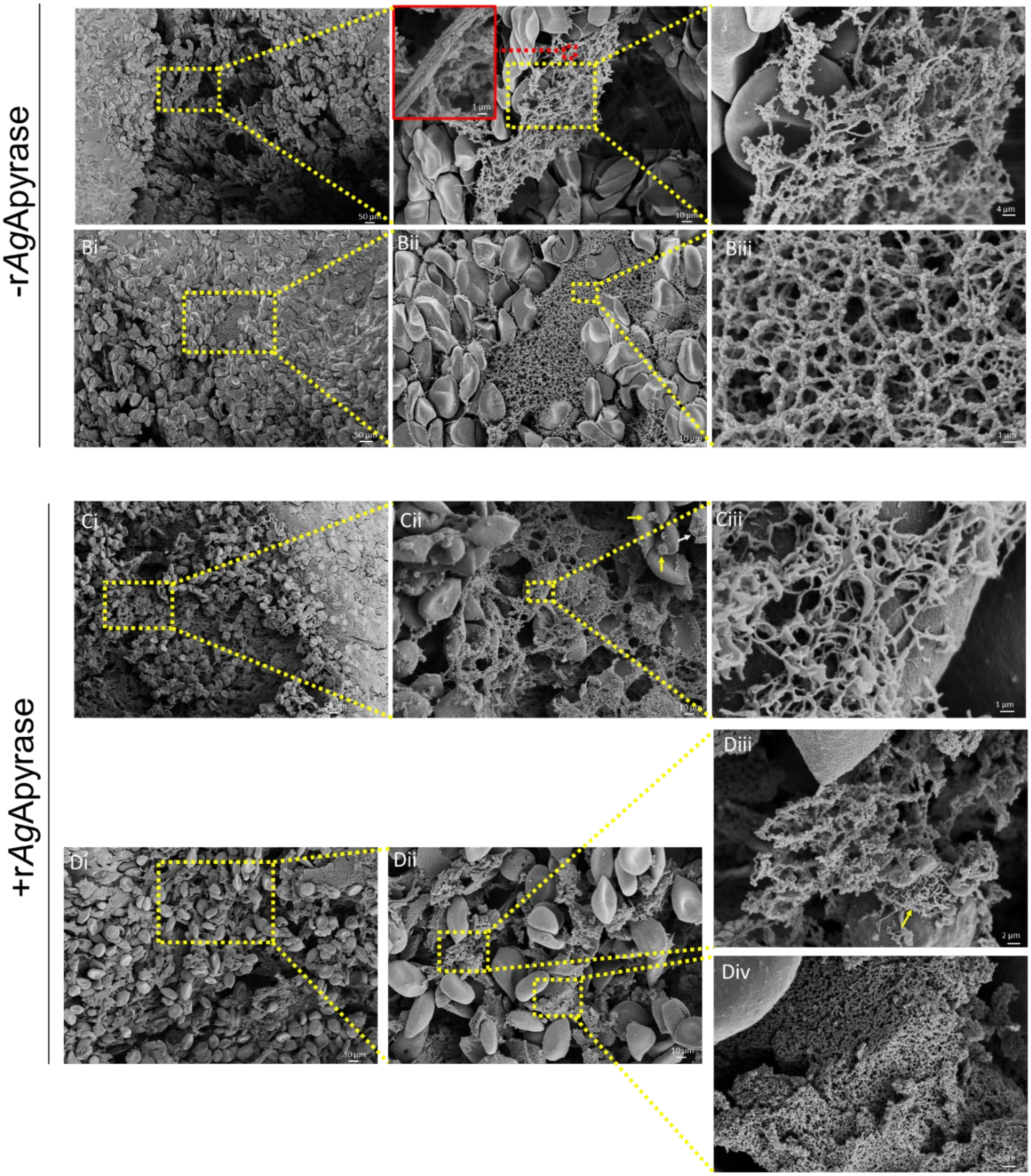


**Fig S7:** Scanning electron microscopy (SEM) of blood boluses before and after supplementation with rAgApyrase. *An. gambiae* female mosquitoes were fed on mice before or after intravenous injection of rAgApyrase. Midguts were dissected at 30 min post feeding. White arrows indicate aggregated platelets, yellow arrows indicate individual platelets.

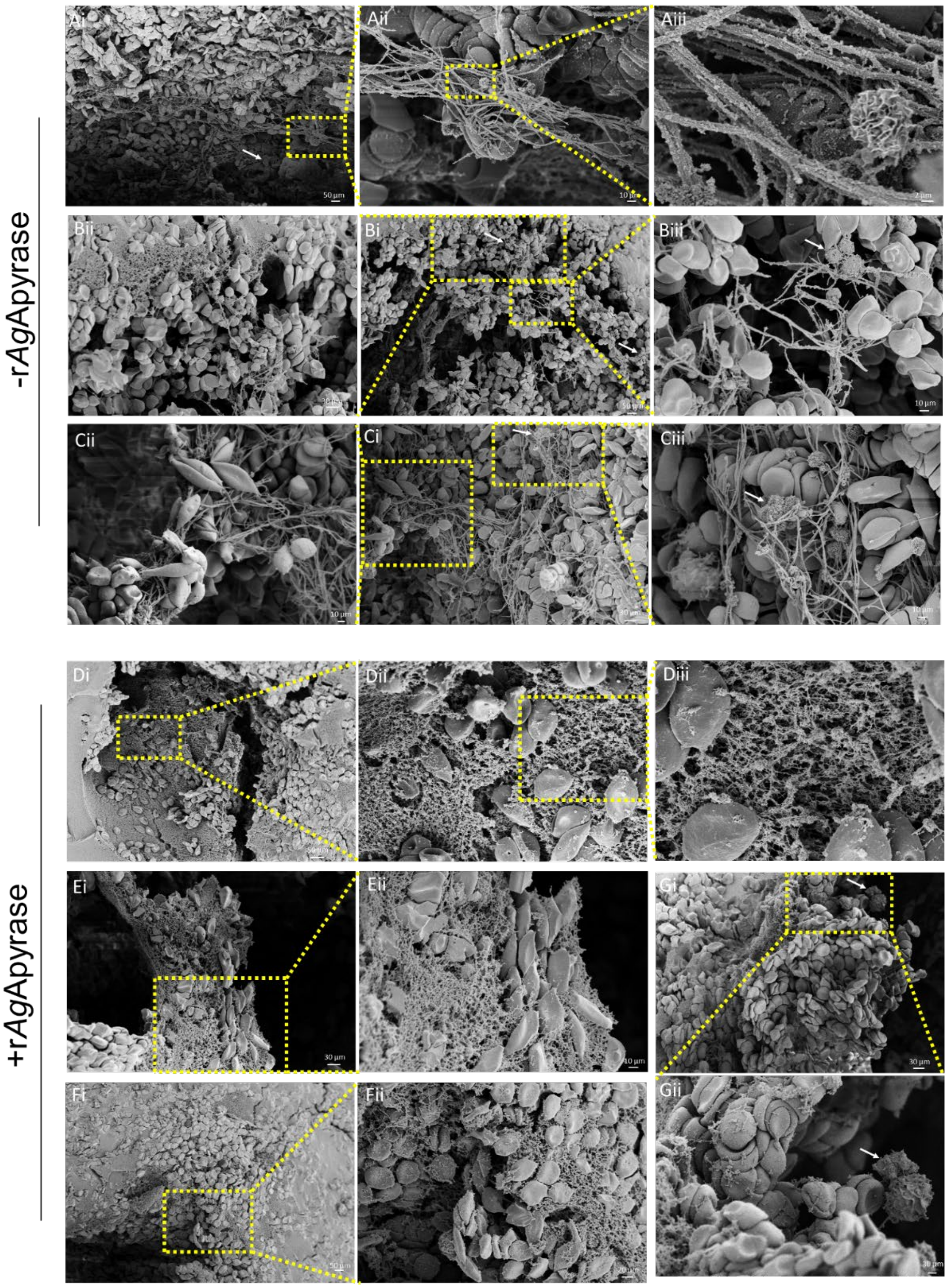


**Fig S8:** Scanning electron microscopy (SEM) of blood boluses before and after supplementation with rAgApyrase. *An. gambiae* female mosquitoes were fed on mice before or after intravenous injection of rAgApyrase. Midguts were dissected at 4 h shown post feeding. White arrows indicate aggregated platelets.

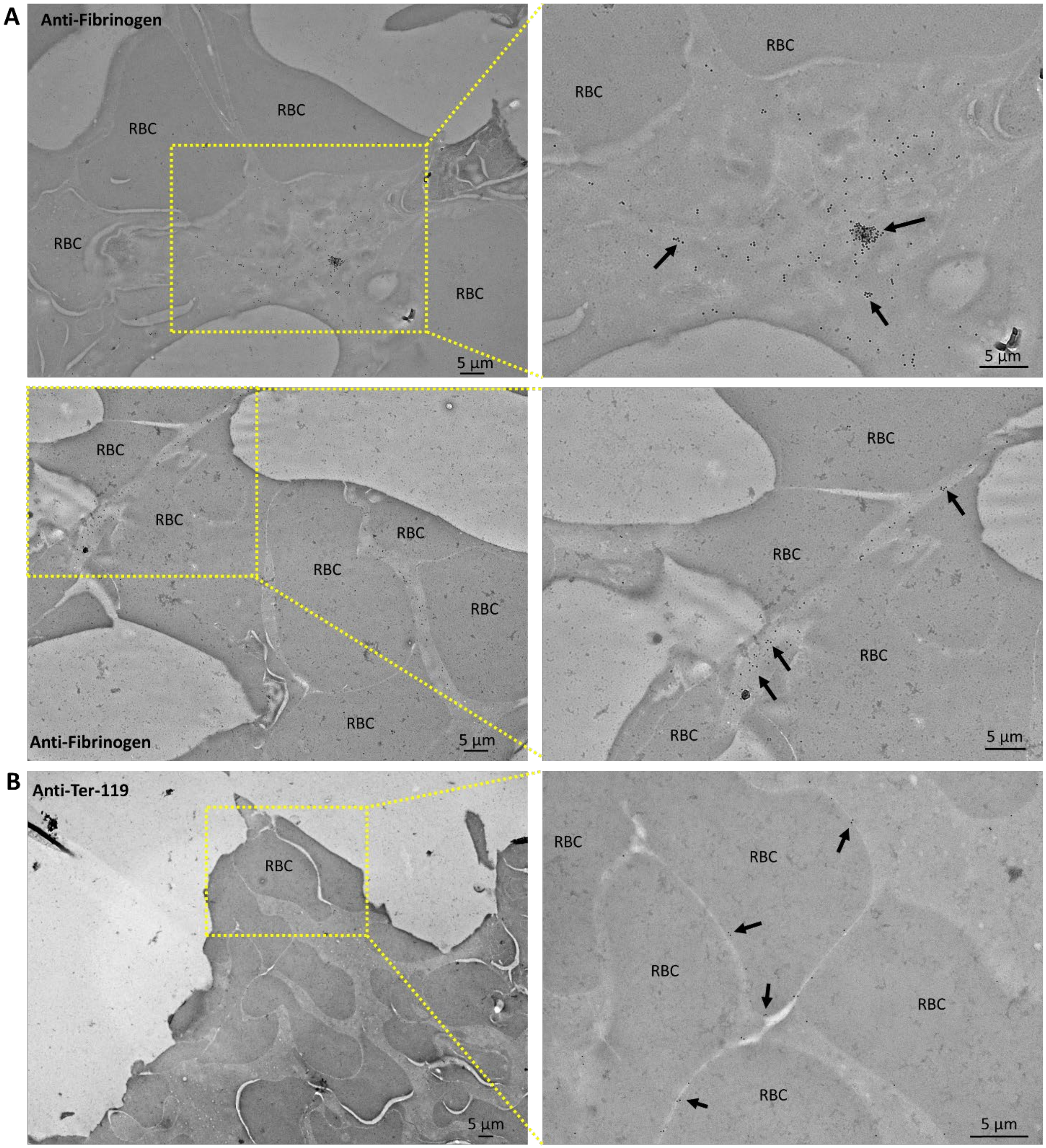


**Fig S9:** Immuno-TEM detecting fibrinogen in the blood bolus. Transmission electron microscopy (TEM) of the blood bolus from mosquito midguts isolated 30 min post blood feeding and stained with anti-fibrinogen or anti-TER-119 antibodies to confirm the presence of fibrinogen/fibrin. In panel A, the arrows point to the gold particles showing the fibrinogen staining in the regions in between the red blood cells (RBC), whereas in panel B, the arrows point to the gold particles showing the staining for the RBC surface protein Ter-119.

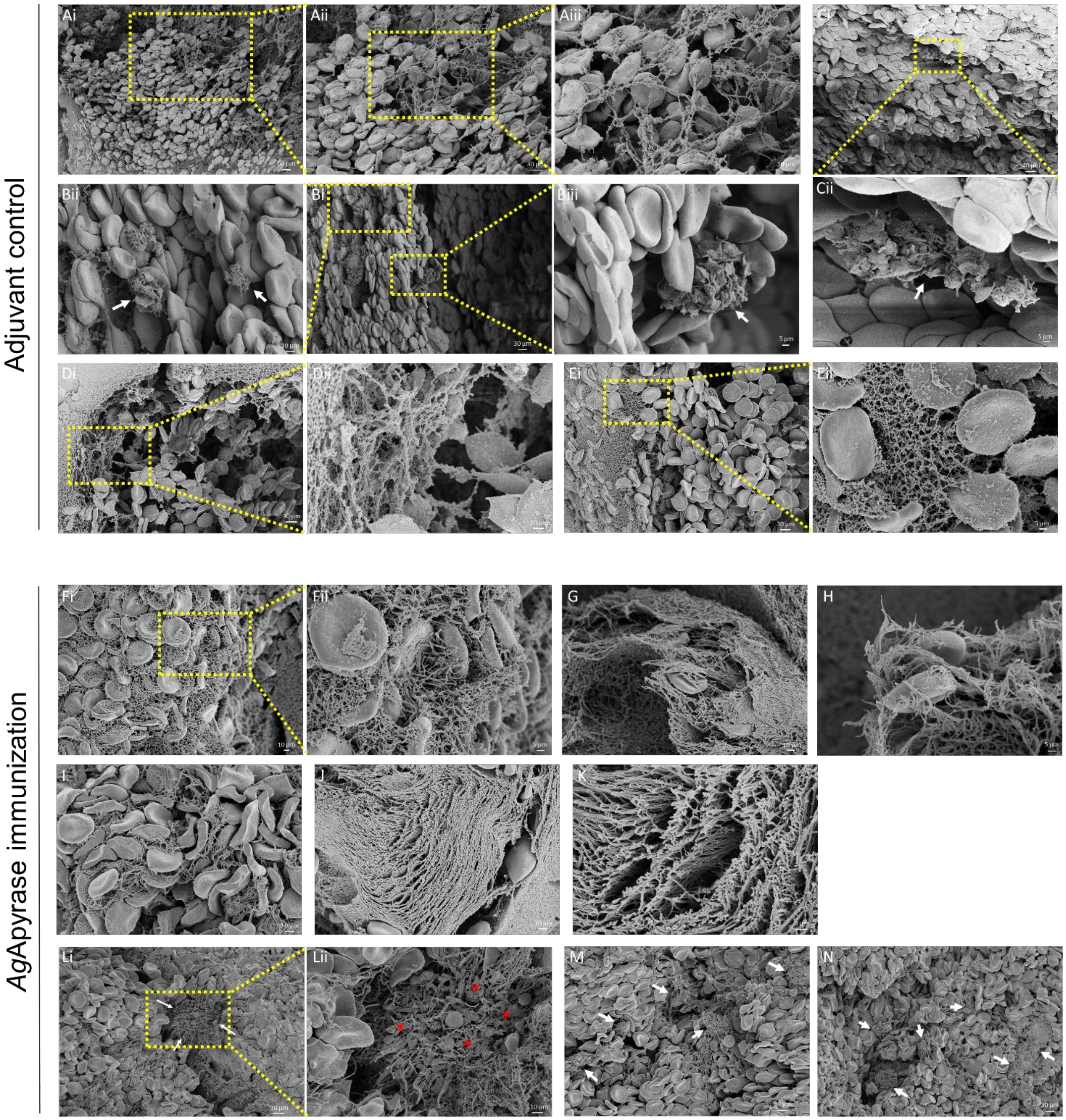


**Fig S10**: SEM of blood boluses from mosquitoes fed on rAgApyrase immunized mice. Control mosquitoes were fed on mice treated with adjuvant. Midguts were dissected 30 min post feeding. White arrows indicate aggregated platelets. Red asterisks show fragmentation of platelets into smaller vesicular bodies.

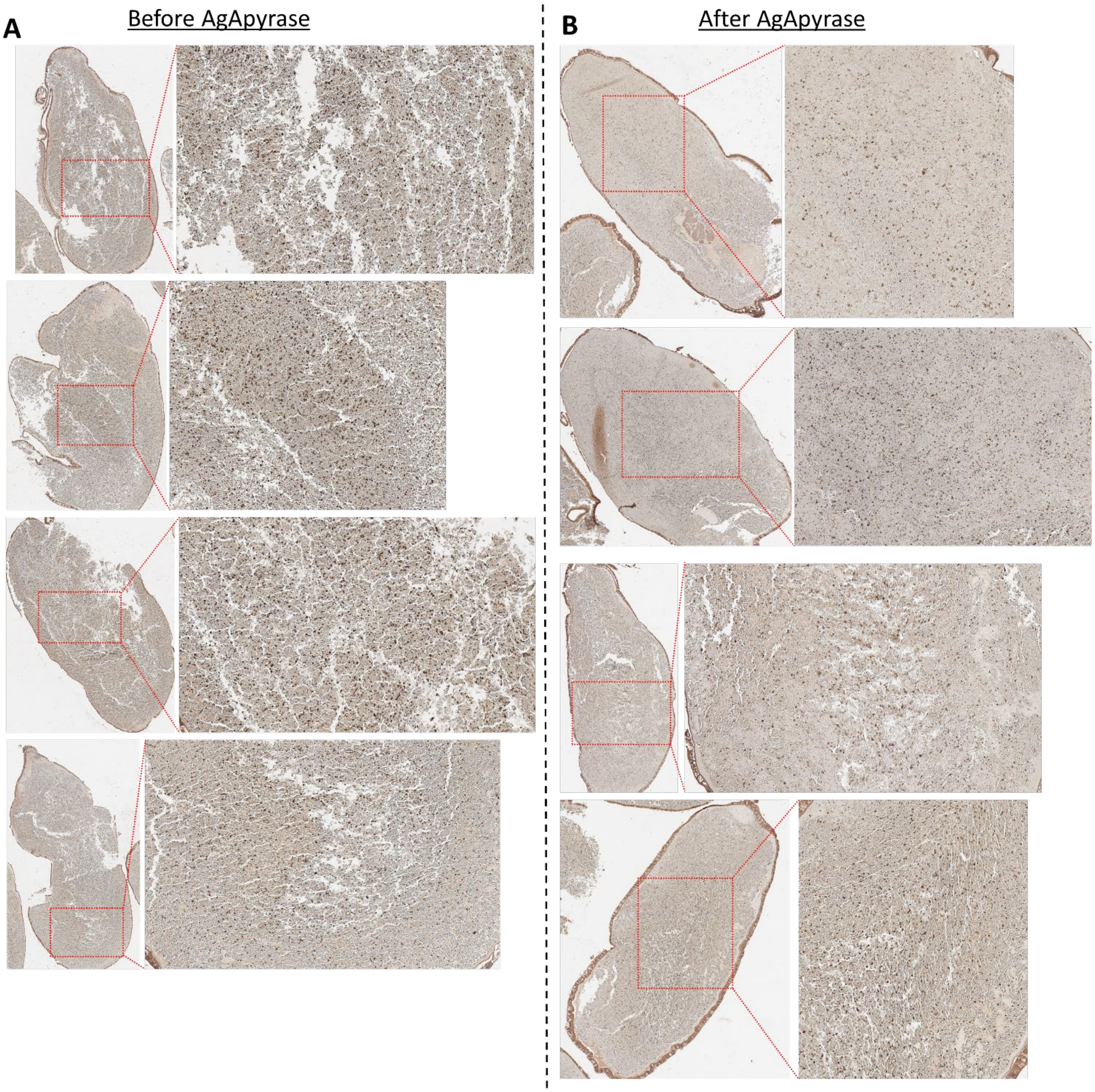


**Fig S11:** Immunohistochemistry of mosquito blood boluses before (A) and after (B) supplementation with rAgApyrase showing platelet activation by staining with P-selectin (dark brown spots). A decrease in staining for the after treatment is observed.

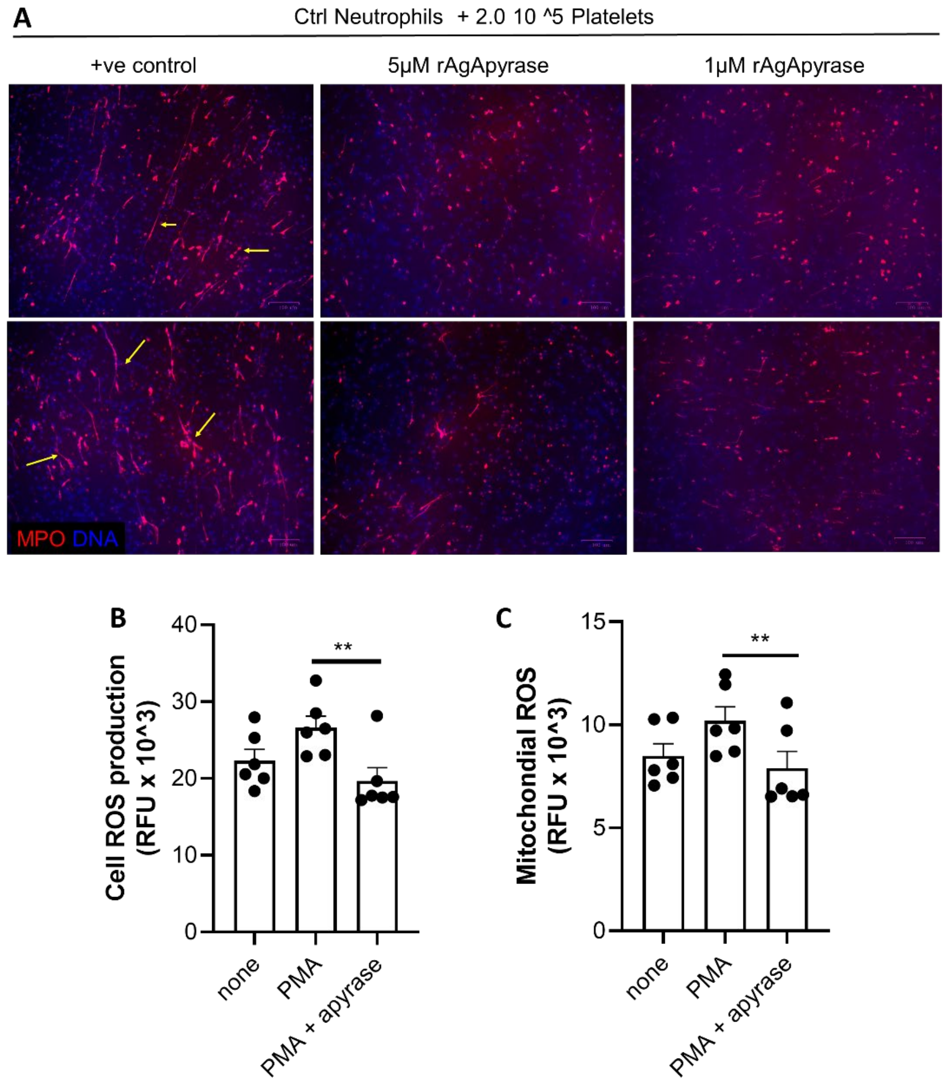


**Fig S12:** AgApyrase inhibits platelet-mediated NET formation and neutrophil ROS production. **(A)** Neutrophils were incubated with platelets in the presence or absence of rAgApyrase. NETs were quantified by immunofluorescence microscopy using an anti-MPO (myeloperoxidase) antibody. DNA was stained with DAPI. Yellow arrows point released neutrophil DNA stained with MPO. Representative images of two independent experiments. (B and C) AgApyrase inhibits neutrophil ROS production. Whole cell (A) or mitochondrial (B) ROS production in neutrophils activated or not with phorbol myristate acetate (PMA) and incubated in the presence or absence of rAgApyrase. One-way ANOVA with Friedman test. **P =0.003.

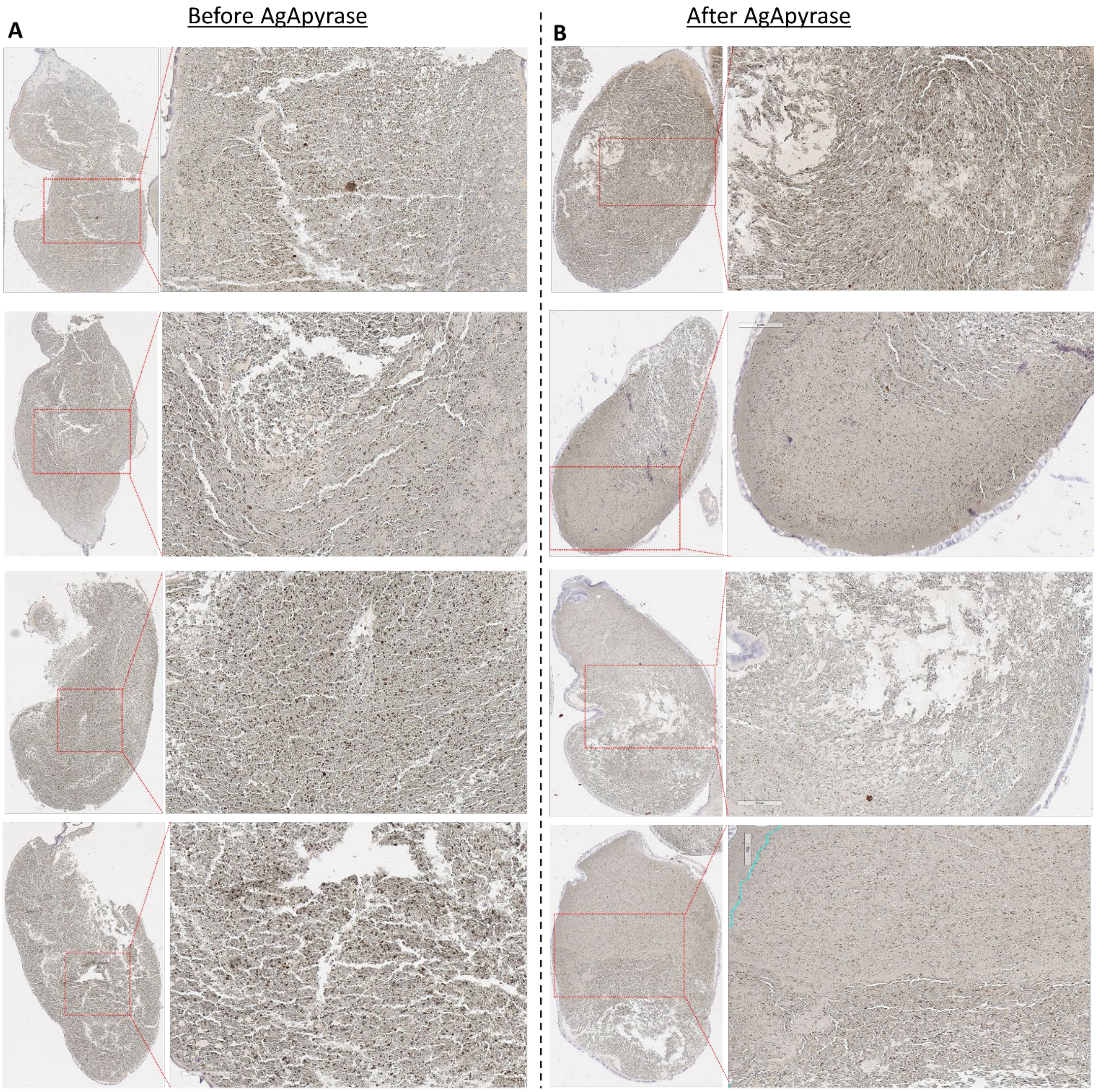


**Fig. S13:** Immunohistochemistry of mosquito blood boluses before (A) and after (B) supplementation with rAgApyrase stained with an anti-neutrophil elastase antibody (dark brown spots).

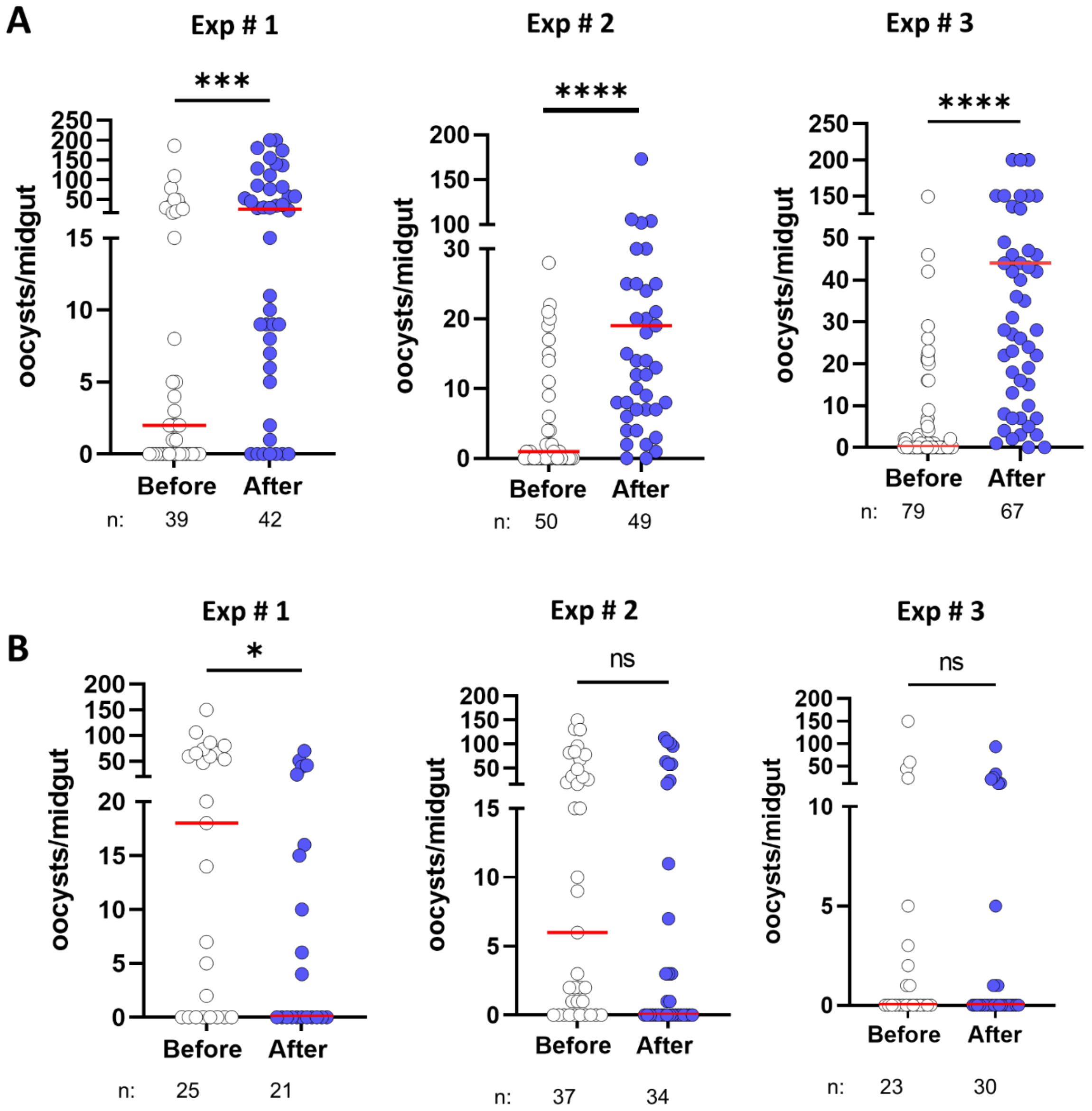


**Fig S14**: Effect of rAgApyrase on *Plasmodium* infection. **(A)** *An. gambiae* mosquitoes were fed on a *P. berghei* infected mouse for 15min (Before group) and the same infected mouse was intravenously supplemented with rAgApyrase. A different group of mosquitoes were fed for 15min (After group) on the injected mouse. Mosquito midguts were dissected 10 days later, and oocysts were counted. AgApyrase supplementation increases *Plasmodium* infection of mosquito midguts. Data from three individual experiments shown as pooled data in Fig. 4A. **(B)** On supplementation of heat denatured rAgApyrase, there is a decrease in the *Plasmodium* infection. Data from three individual experiments shown as pooled data in Fig. 4A. n= no. of mosquitoes/ experiment. Horizontal red line represents the median oocyst number per mosquito, each dot represents a mosquito. Groups were compared with two-tailed t-test followed by Mann-Whitney comparison test. ****P <0.0001; ***P <0.0006; *P<0.045.

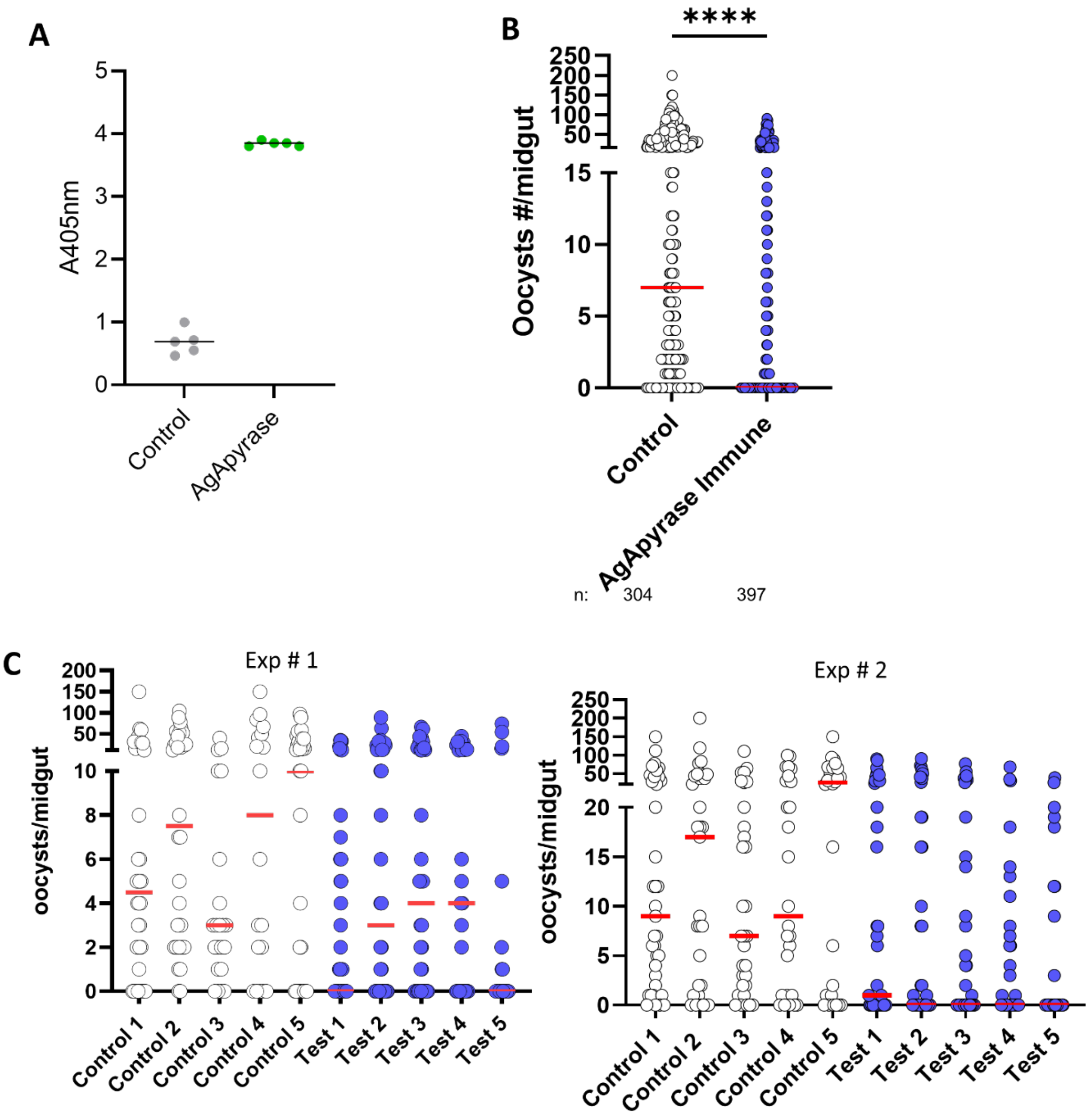


**Fig S15.** AgApyrase immunization inhibits *Plasmodium* midgut infection. **(A)** BALB/c mice were immunized with recombinant AgApyrase using Magic Mouse adjuvant. Adjuvant only was used as a control for the experiment. The antibody titers were determined using ELISA. Each dot represents data from one animal. **(B)** The control and immunized mice were infected with *P. berghei* and *An. gambiae* mosquitoes were fed for 15min. Mosquito midguts were dissected 10 days later, and oocysts were counted. AgApyrase significantly inhibits *Plasmodium* midgut infection. Pooled data for no. of oocysts/ midgut fed on 10 individual animals shown. **(C)** Data from mosquitoes fed on each individual animal of the data shown in panel B. Data obtained from two independent immunization experiments (experiment #1 and #2). Control: Adjuvant only; Test: Apyrase Immunized. n= no. of mosquitoes.

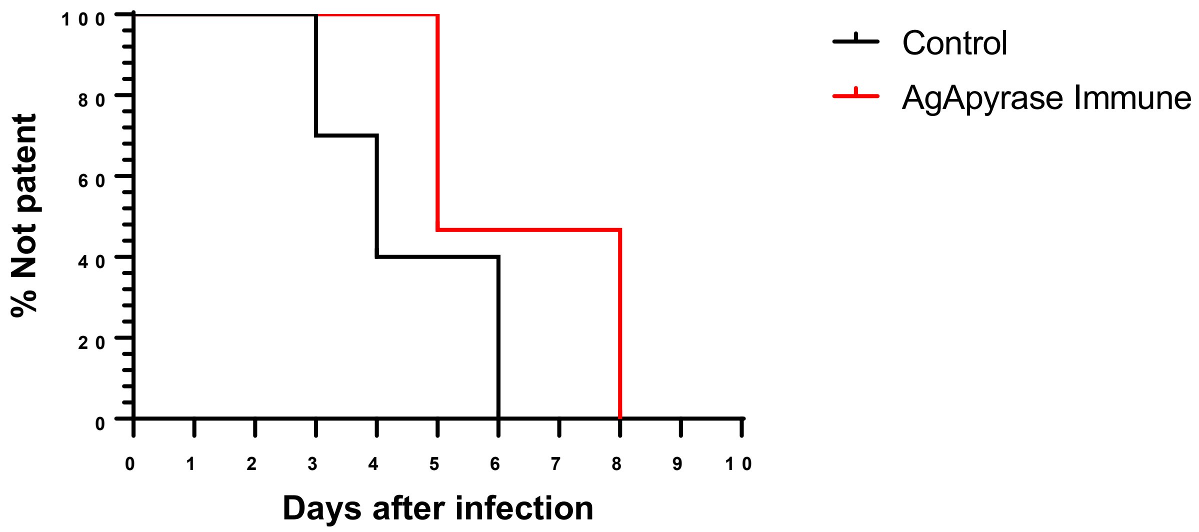


**Fig S16.** Pre-patency for control and apyrase immunized mice. Sporozoite infectivity was determined by the day of appearance of blood-stage parasites in peripheral blood (patency) by Giemsa staining. Data pooled from two independent experiments shown in Fig. 4D and 4E.

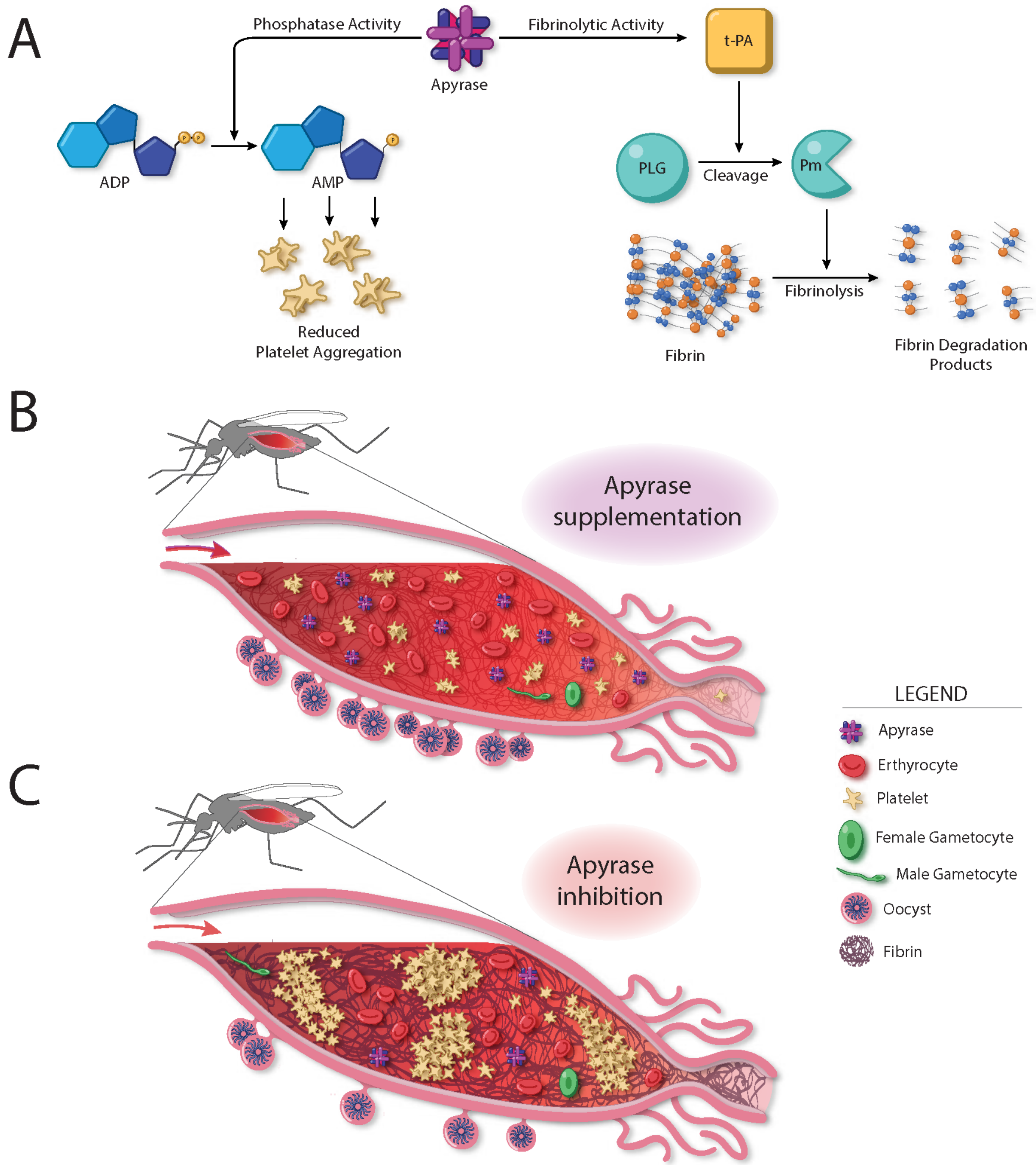


**Fig S17:** Proposed model for the role of AgApyrase in the mosquito midgut. (A) AgApyrase acts as a classical apyrase with phosphatase activity by hydrolyzing ADP to release AMP and phosphate and therefore, prevents ADP-mediated platelet aggregation. AgApyrase activates tissue plasminogen activator (tPA) which in turn activates plasminogen to plasmin. Plasmin enhances the degradation of fibrin. **(B)** Fibrin polymerization is detected in the mosquito midgut within minutes of an infectious blood meal ingestion. The salivary apyrase ingested during blood feeding, enhances fibrin degradation, and inhibits platelet aggregation thus facilitating the migration of *Plasmodium* gametes in the blood bolus and promoting parasite infection. **(C)** Inhibition of apyrase with anti-apyrase antibodies results in the formation of a denser fibrin network and increased platelet aggregation, which interferes with *Plasmodium* gamete migration and parasite infectivity.
